# Structural dynamics of SARS-CoV-2 nucleocapsid protein induced by RNA binding

**DOI:** 10.1101/2021.08.27.457964

**Authors:** Helder Veras Ribeiro Filho, Gabriel Ernesto Jara, Fernanda Aparecida Heleno Batista, Gabriel Ravanhani Schleder, Celisa Caldana Tonoli, Adriana Santos Soprano, Samuel Leite Guimarães, Antonio Carlos Borges, Alexandre Cassago, Marcio Chaim Bajgelman, Rafael Elias Marques, Daniela Barreto Barbosa Trivella, Kleber Gomes Franchini, Ana Carolina Migliorini Figueira, Celso Eduardo Benedetti, Paulo Sergio Lopes de Oliveira

## Abstract

The nucleocapsid (N) protein of the SARS-CoV-2 virus, the causal agent of COVID-19, is a multifunction phosphoprotein that plays critical roles in the virus life cycle, including transcription and packaging of the viral RNA. To play such diverse roles, the N protein has two globular RNA-binding modules, the N-(NTD) and C-terminal (CTD) domains, which are connected by an intrinsically disordered region. Despite the wealth of structural data available for the isolated NTD and CTD, how these domains are arranged in the full-length protein and how the oligomerization of N influences its RNA-binding activity remains largely unclear. Herein, using experimental data from electron microscopy and biochemical/biophysical techniques combined with molecular modeling and molecular dynamics simulations, we showed that, in the absence of RNA, the N protein formed structurally dynamic dimers, with the NTD and CTD arranged in extended conformations. However, in the presence of RNA, the N protein assumed a more compact conformation where the NTD and CTD are packed together. We also provided an octameric model for the full-length N bound to RNA that was consistent with electron microscopy images of the N protein in the presence of RNA. Together, our results shed new light on the dynamics and higher-order oligomeric structure of this versatile protein.

## Introduction

All coronaviruses, including the Severe Acute Respiratory Syndrome Coronavirus 2 (SARS-CoV-2), the causative agent of the Coronavirus disease 2019 (COVID-19) pandemics, possess an organized nucleocapsid formed by a ribonucleoprotein (RNP) complex surrounded by a lipid envelope [1–3]. The major component of the RNP complex is the nucleocapsid (N) protein, one of the four structural proteins of coronaviruses and also the most abundantly expressed viral protein in infected host cells [2].

N proteins are conserved among coronaviruses and are known to play multiple roles in the virus life cycle [4]. In addition to packaging the viral genomic RNA, N proteins are required for genome replication, transcription and translation, and for the assembly of the RNPs into newly formed viral particles [5–13]. This functional diversity is intimately linked to the dynamic structure of the N protein and its ability to bind and alter the RNA structure [9,14].

Coronaviruses N proteins are composed of two structured and globular domains represented by the N-(NTD) and C-terminal (CTD) domains, both of which are capable of binding single-stranded RNA and DNA molecules [15–24]. The NTD has an extensive basic U-shaped RNA-binding cleft implicated in the binding and melting of transcription regulatory sequences (TRS) needed for transcription of sub-genomic RNAs [9,14]. The CTD is responsible for the protein dimerization and it also forms a positively charged groove thought to contribute to the recognition of the packaging signal (PS) and to the assembly of the RNP into the virion particle [13,17,23]. In addition to the CTD, the flexible C-terminal tail also seems to influence protein oligomerization by promoting protein tetramerization [25–27]

The NTD and CTD are connected by a central disordered serine and arginine-rich region, denoted as SR linker. This linker region is also proposed to play fundamental roles in protein oligomerization and function. Of note, the SR linker was shown to be modified by phosphorylation, which not only reduces the affinity of the protein for the RNA, but also drives a liquid phase separation of the N protein with the RNA and other virus proteins and host cell components [28–30]. Recently, mutations on SR-linker have been associated with the emergence of the high-transmissibility SARS-CoV-2 lineage B.1.1.7 [31], which makes vital not only the determination of NTD or CTD structures, but also the understanding of the full organization of N protein with its disorder linkers and tails.

Due to its dynamic structure and multiple oligomeric organization depending on environmental conditions, and to the fact that the N protein is also modified by phosphorylation [28,29,32], no three-dimensional (3D) structures are available for coronaviruses full-length N proteins. Here, by combining electron microscopy and biophysical experimental analysis with molecular modeling and molecular dynamics simulations, we propose structural models for the full-length SARS-CoV-2 N protein in the absence and presence of RNA. These models not only support current experimental data, but also provide a framework for understanding the multifunctional role of the N protein.

## Results

### Dimers of full-length N adopts an extended conformation in the absence of RNA

In order to shed new light on the structure of full-length N and to understand how the NTD and CTD are oriented to each other, the recombinant N protein produced in *E. coli* was purified by affinity and size-exclusion chromatography (SEC) and analyzed by negative stain electron microscopy (NSEM). We noticed that the N protein appeared as a single peak in the SEC and as a single band in denaturing gel electrophoresis (Figure 1A and 1B). Further, N protein contained traces of bacterial RNA in non-denaturing gels, evidenced by the 260/280 nm absorption ratio about 1.8 for this N protein sample. Accordingly, treatments with RNase A, but not DNase I, removed most of the nucleic acid associated with the protein (Figure 1B), reducing the 260/280 nm absorption ratio to ~0.68. Notably, removal of the contaminating RNA with RNase A led to a change in the oligomeric state of the N protein (Figure 1B).

**Figure 1.**
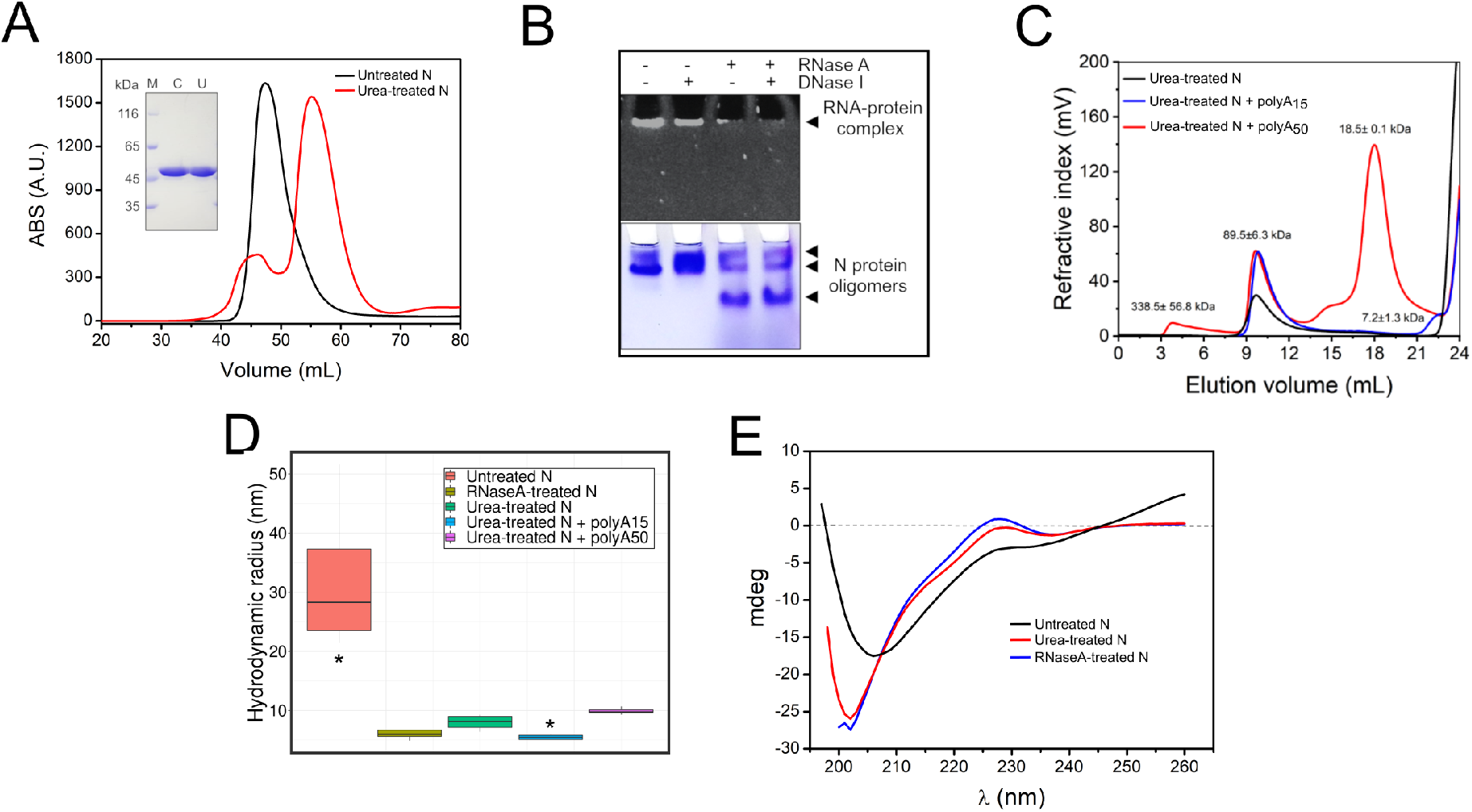
The N protein forms high molecular-weight complexes with RNA but is a dimer in solution in the absence of RNA. A- SEC showing that the untreated N protein sample (black line) elutes as a single peak in the void volume of the gel filtration column whereas the urea-treated sample (red line) elutes as a major peak of a higher elution volume. The purity of these proteins samples was evaluated by SDS-PAGE, which shows single bands of the expected size for the recombinant N protein (inset). For urea-treated samples, all experimental analysis were carried out using fractions corresponding to the center of the major peak, since the minor peak likely corresponds to the protein still carrying nucleic acid. B- Native gel electrophoresis stained with ethidium bromide (upper panel) or Coomassie blue (lower panel) showing that the N protein forms high molecular-weight complexes with bacterial RNA, as treatments with RNase A, but not DNase I, dissolve these complexes (arrowhead). The RNase A treatment also changed the oligomeric forms of N protein (arrowheads) in solution (lower panel). C- SEC-MALS analysis of urea-treated N protein in the absence or presence of polyA15 or polyA50. The chromatogram shows that the N protein treated with urea as well as in the presence of polyA15 are dimers in solution, with a mean molecular mass of 89.5 ± 6.3 kDa. In addition to the peak of dimers, urea-treated N presents a higher molecular-weight peak (338.5 ± 56.8 kDa) and a lower molecular-weight peak (18.5 ± 0.1 kDa) D- DLS measurements showing the hydrodynamic radius of untreated N protein carrying bacterial RNA (red boxplot), N protein treated with RNase A (brown boxplot) and N protein treated with urea in the absence (green box plot) or presence of polyA15 (blue boxplot) or polyA50 RNA (purple boxplot). Statistical comparison was performed using Dunnett-Tukey-Kramer pairwise multiple comparison test adjusted for unequal variances and unequal sample sizes and P value was estimated from the confidence interval (see methods for details). *P<0.05 comparing urea-treated N with the other conditions. E- CD plot showing that CD curves of N protein samples treated with RNase A (blue) and urea (red) are compatible These curves are more similar to each other than the one of N protein samples carrying bacterial RNA (black).

To investigate how the N protein behaves in the absence of RNA, the recombinant protein was treated with urea and high salt concentration to remove bound RNA [28,33,34], and purified by affinity chromatography and SEC. The urea-treated N, which presented a 260/280 nm absorption ratio of ~0.5, eluted as a major peak with a higher elution volume compared with the untreated protein (Figure 1A). This sample showed a molecular mass of 89,5 ± 6.3 kDa and hydrodynamic radius of 8.0 ± 1.3 nm determined by SEC-MALS and DLS, respectively, consistent with a dimer in solution (Figure 1, C and D). Noteworthy, the hydrodynamic radius is close to the one recently reported for N protein purified under similar conditions [35]. Also, these results are in line with literature data showing that N protein readily oligomerizes into dimers [36,37]. The urea-treated N showed a circular dichroism profile comparable to that of the protein treated with RNase A (Figure 1E), indicating that the urea treatment did not substantially alter the protein secondary structure. Both curves are close to previous reported N protein CD profiles [37]. On the other hand, CD profile of untreated N protein presents a positive ellipticity at 250-260 nm which represents the contribution of the nucleic acids present in this sample, most likely RNA,

Negative stain images produced from the urea-treated N samples showed various rounded particles with dimensions smaller than 5 nm (Figure 2A). Thus, the dimensions of these particles are not directly correlated with the hydrodynamic radius determined experimentally for the full-length N protein (Figure 1D). Nevertheless, these 5 nm-particles can represent the globular domains of the N protein in an extended conformation, in which each particle corresponds to individual NTD or CTD units. N protein disorder regions, such as SR-linkers and tails, do not present sufficient contrast to be distinguished from the background noise in this NSEM.

**Figure 2.**
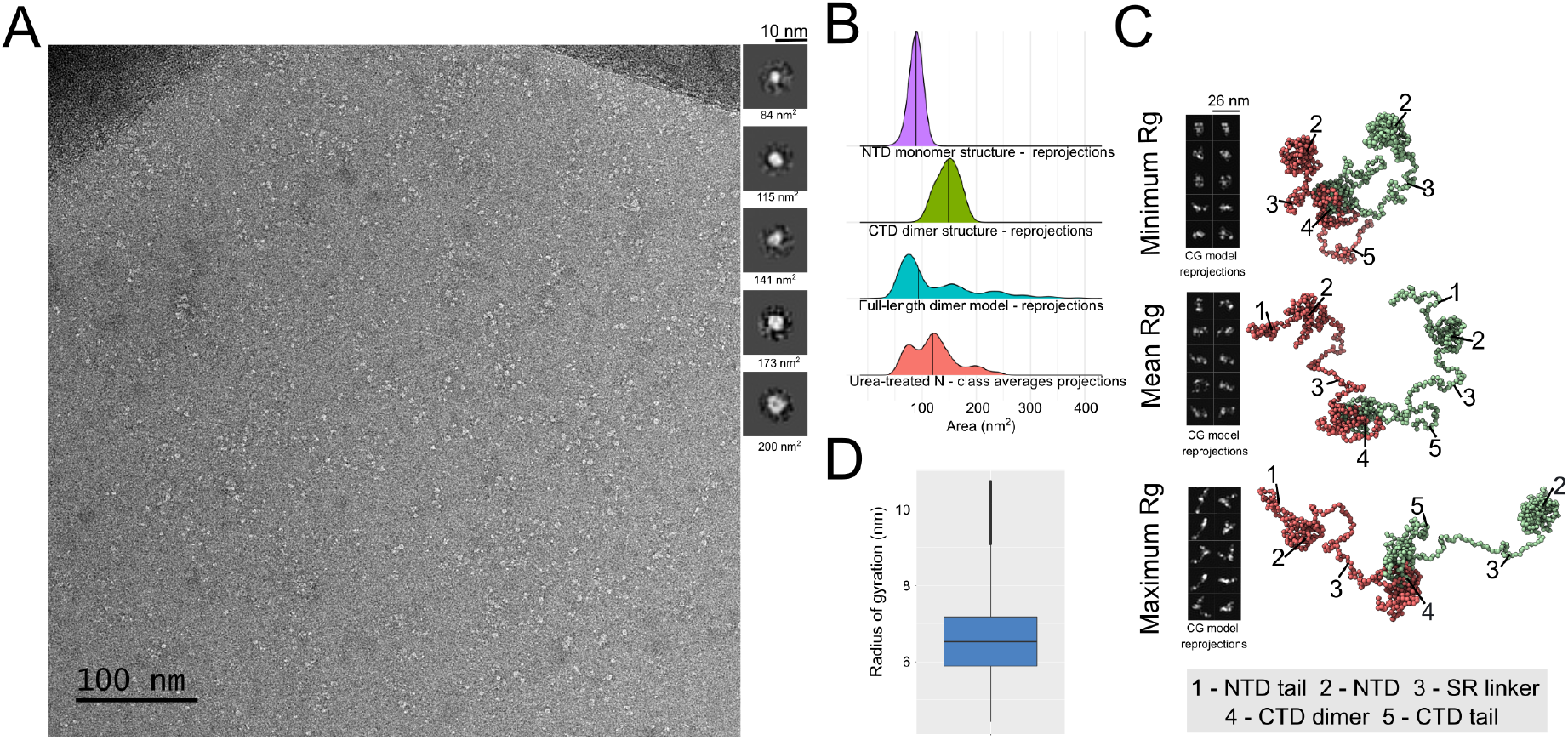
Dimers of full-length N show an extended conformation in the absence of RNA. A- Representative negative stain image of urea-treated N in the absence of RNA. Insets on the right present representative class averages from a total of 100 classes produced by the 2D classification of 3500 particles. The area corresponding to each particle is presented. B- Density plots of area distribution estimated for simulated 2D reprojections from NTD monomer structure (PDB ID: 6M3M), CTD dimer structure (PDB ID: 6WZO) or full-length dimer reprojections, or for class averages of urea-treated N. In the case of full-length dimer reprojections, when N protein structured domains (NTD or CTD) were presented as separated density regions in the reprojections, the area of these regions were computed separately. For this reason, we can see three main peaks in full-length N dimer distribution: the first peak corresponding to isolated NTD domains, the second to isolated CTD domains and the third to combinations of these domains close to each one in the 2D reprojected image (see methods for details). C- Representative frames of the minimum (upper panel), mean (middle panel) or maximum (lower panel) radius of gyration obtained from the CG simulations of the full-length N protein dimer. 2D simulated reprojections from each frame that were used to estimate area are presented at the left. N protein domains are numbered. D- Boxplot of the radius of gyration estimated from five independent CG simulations of full-length N protein dimer in absence of RNA.

To investigate this hypothesis, we picked 3500 particles and performed a two-dimensional (2D) classification into 100 classes to calculate the area of each class average (Figure 2A, inset). We compared the distribution of areas of the urea-treated N class averages with the distribution of areas of simulated reprojections derived from the monomeric NTD and dimeric CTD atomic structures (Figure 2B) (see methods and Figure S2B for details). Notably, we found that the area distributions of the urea-treated N protein fitted well into the area distributions of the NTD monomer and CTD dimer (Figure 2B). This suggested that the particles observed in the NSEM images (Figure 2A) likely correspond to isolated NTD and CTD globular domains of the N protein dimer in extended conformations. The idea that N protein adopts an extended conformation in absence of RNA is supported by coarse-grained (CG) molecular dynamics simulations of the full-length N protein dimer, which show that the NTD regions move freely in relation to the CTD dimer, producing a variety of conformers (Figure 2C) with an estimated mean radius of gyration of 6.6 nm (Figure 2D). This value is close to the radius of gyration of a SARS-CoV-1 N protein construction without N- and C-terminal tails (6.1nm), determined by SAXS [19]. The Rg calculated from CG simulations is numerically different from the experimentally estimated Rh described above, which is expected since they represent two different physical properties [38,39]. Thus, in CG simulation analysis, we used the Rg as a metric to estimate the compaction upon RNA binding, while the values of Rh were used for comparing compaction in DLS experiments.

The CG simulations do not point to NTD-CTD contacts (Figure S1), supporting the idea that the large density of positive charges on the surface of the NTD and CTD causes repulsion of these domains, which is favored by the flexibility of the SR linker. The simulated 2D reprojections derived from representative 3D models (with minimum, mean or maximum radius of gyration) of the CG simulation also illustrate the diversity of N protein conformations. These simulated 2D reprojections show density regions of similar size of the particles observed in the NSEM images (Figure 2, A and C). In the case of full-length N dimer conformations obtained from CG, depending on its orientation, the reprojections of these conformations can be seen as three high density points that correspond to the structured domains (two NTD and one CTD dimer) in an extended conformation (Figure 2C and Figure S2B), producing the multimodal distribution observed in Figure 1B. The area distributions of the of the urea-treated N class averages overlap with the area distributions of the simulated reprojections of the full-length N dimer (Figure 2B). Together, our data suggest that dimers of full-length N show extended and protein highly flexible conformations in the absence of RNA.

### Distinct forms of the N protein bound to RNA were revealed by NSEM

Because no 3D structures of full-length N in complex with RNA are available and only few microscopy studies have tried to report the structural organization of N protein [28], we inspected the N protein samples not treated with urea or RNase A by NSEM. The NSEM micrographs presented a very informative panel of N protein in the presence of nucleic acid and showed that untreated N protein preparations contained a myriad of particles and aggregates that are consistent in size with the DLS measurements (Figure 1D and Figure 3).

**Figure 3.**
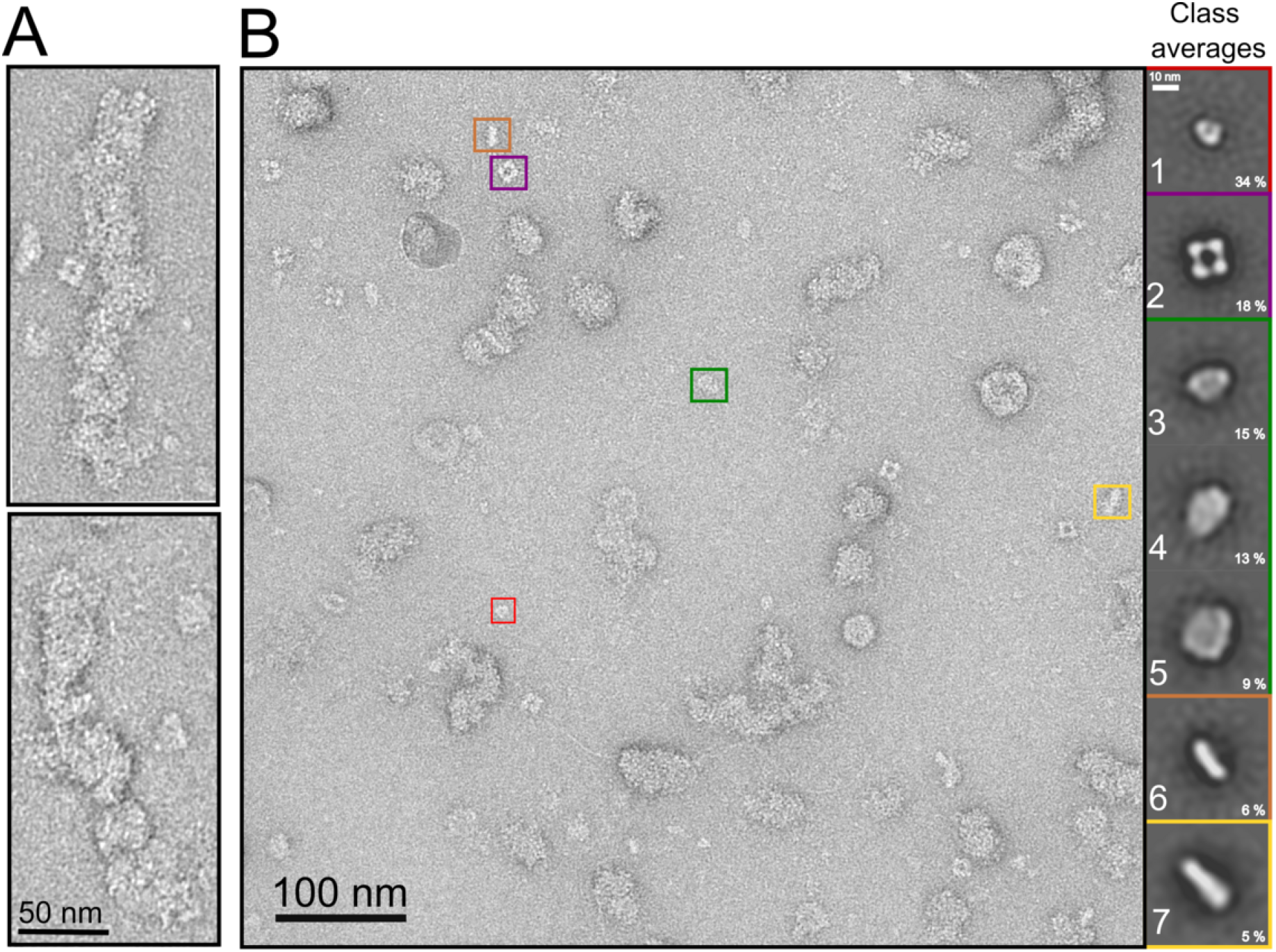
Representative negative stain micrographs of untreated N protein samples containing bacterial RNA. A- Detail of large helical-like structures with 20 – 30 nm width. B- Representative micrograph showing a wide range of particles and aggregates. Smaller particles, with less than 20 nm in diameter are shown (colored boxes). In insets, class averages (1 to 7) of picked particles with sizes up to 20 nm in width from one hundred electron micrographs: class-1 (toroid-like particles), class-2 (square-like particles), classes 3 to 5 (elliptical and round-like particles), and classes 6 and 7 (rod-like particles). The percentage of each class average in relation to all picked particles are indicated.

The larger aggregates with a width of 20 – 30 nm (Figure 3A) resemble the helical structures of the linearly arranged N protein isolated from the transmissible gastroenteritis (TGEV) coronavirus [40]. Smaller particles with less than 20 nm in diameter were also observed (Figure 3B, colored boxes). Because these particles were quite abundant and structurally diverse, and showed dimensions consistent with N protein particles isolated from SARS-CoV-2 and related virus [41–43], we classified them according to their size and shape (Figure 3C).

The 2D class averages analysis revealed seven particle classes (Figure 3B). Class-1 (toroid-like) particles comprised rounded particles with a weak central density. These particles, which were the smallest and most compact (between 7 and 10 nm wide), resemble those corresponding to the mouse hepatitis virus (MHV) N protein dimer [41,42].

Class-2 particles comprised square-like particles approximately 14 nm wide. Notably, these particles have not yet been associated with the SARS-CoV-2 N protein; however, they resemble the RNPs isolated from the human coronavirus 229E (HCV229E) and MHV3[44]. The class-2 particles also resemble those of the M1 influenza A protein and Schmallenberg orthobunyavirus RNP, where each globular unit located at the square vertices corresponded to a single structured protein domain [45,46].

Class averages 3 to 5 comprised elliptical and rounded particles ranging from 11 to 19 nm in length, whereas class averages 6 and 7 comprised rod-like structures of 16-20 nm in length (Figure 3B).

Although the diversity of these particles hinders the use of classical single-particle approaches needed for 3D structure high-resolution, we used the 2D class average analysis to estimate particle dimensions and to guide molecular modeling and dynamics simulations to gain insights into the conformational states of full-length N bound to RNA. We focused our analysis on the most abundant particles represented by classes 1 and 2, which are also the particles that showed well-defined features and dimensions (Figure 3B).

### N protein dimers adopt a packed conformation in the presence of RNA

As reported recently [36,37] and shown in Figure 1C, the N protein devoid of RNA is a dimer in solution, which was consistent with the notion that dimers are the fundamental oligomeric unit of full-length N. The toroid-like particles observed in Figure 3B had dimensions ranging from 7 to 10 nm, which were smaller than the expected N protein diameter estimated from Rh (2 × 8.0 nm in diameter) or Rg (2 × 6.6 nm in diameter) as described above. Thus, we reasoned that these particles could represent N protein dimers in a more compact conformation due to the presence of RNA. This idea was supported by the observation that the RNase A-treated N, which may still have RNA bound to it, exhibited a smaller hydrodynamic radius (5,9 ± 0,7 nm) compared with the urea-treated protein. (Figure 1D).

To investigate this hypothesis, the urea-treated N was incubated with a polyA15 or polyA50 RNA and the hydrodynamic radius was determined by DLS. We found that the N protein incubated with polyA15, but not polyA50, showed a significant reduction in the hydrodynamic radius (5.5 ± 0.4 nm), compared with the RNA-free N protein (Figure 1D). Notably, the hydrodynamic radius observed for the protein in the presence of polyA15 was comparable to that of the RNase A-treated protein, which still retains RNA (Figure 1D). On the other hand, the presence of polyA50 RNA increased the hydrodynamic radius of N protein in comparison to the presence of polyA15, producing a hydrodynamic radius slightly higher than the RNA-free N protein (Figure 1D). Interestingly, SEC-MALS analysis of the urea-treated N protein incubated with the polyA50 showed, in addition to the N dimer peak, a small but detectable peak of approximately 340 kDa, which would be consistent with an N protein octamer (Figure 1F). Such peak was not observed with polyA15. In addition, when inspected by NSEM, N protein samples incubated with polyA50 showed larger particles of irregular shape in comparison to samples with polyA15 (Figure S2A), which would infer a long RNA could join two or more dimers of N. These results are in line with the increase in hydrodynamic radius produced by polyA50 and thus suggest that the N protein undergoes protein compaction and possibly oligomerization in the presence of RNA.

To understand how the RNA could drive protein compaction, we performed CG dynamics simulations of dimeric full-length N, in which protein-RNA contacts are mostly driven by electrostatic forces. The simulations were carried out in the presence of single-strand RNA molecules ranging from 10 to 70 nucleotides (nt) in length (Figure 4). We observed that while the CTD dimer, which has the highest density of positive charges, readily associates with the shortest RNA molecules (10 - 20 nt), the NTD units move freely, with no contact with RNA (Figure 4C). Thus, a single short RNA molecule was not able to interact simultaneously with NTD and CTD, and since NTD units moved freely, the distribution of radius of gyration, in this case, is close to the one without RNA (Figure 4B). Two main RNA contact regions can be observed in CTD: the first one includes residues 240 to 280, which encompasses a region with high density of positively charged residues, whereas the second one has no significant number of positively charged residues, but is spatially positioned close to the first contact region. Additionally, an RNA contact region that includes the CTD tail (residues 370 to 390) was also observed. From 30 nt, the single strand RNA is long enough to simultaneously interact to CTD dimer and at least to one NTD unit (Figure 4A and 4C). The RNA contacts in NTD mainly included the region of residues 90 to 110 in N protein and also residues 30 to 50 that were part of NTD tail. From 40 nt, the frequency of RNA contacts with NTD is increased as well as with the SR linker (residues 180 to 200) (Figure 4C), which seems the cause of the shift to smaller radius of gyration observed in the distribution (Figure 4A). As the length of the RNA molecules increases (50 to 60-nt), a single RNA molecule can interact simultaneously with the two NTD units and one CTD dimer bringing these domains together. Full compaction of the N protein dimer was observed with the 40 to 60-nt RNA models with a minimum radius of gyration of 3.7 nm for these models (Figure 4B). However, the presence of multiple RNA binding sites along the N protein could account for the compaction observed experimentally even with shorter RNA such as the polyA15 RNA (Figure 1D). Indeed, CG simulations of the N protein dimer with multiple RNA molecules of 15 nt also presented a reduction in the protein radius of gyration relative to the RNA-free protein (Figure S3), which was thus in line with the DLS results (Figure 1D).

**Figure 4.**
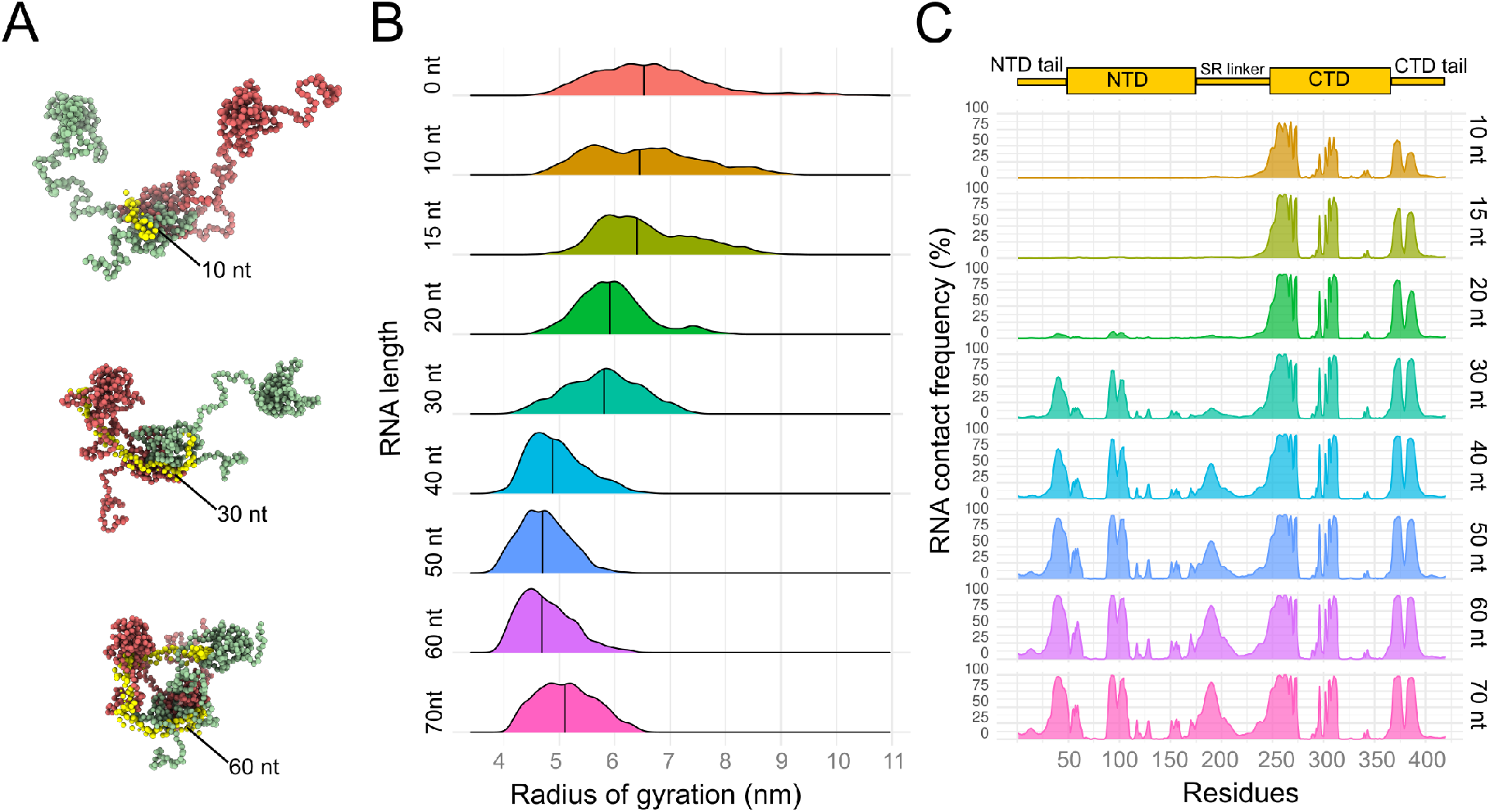
CG dynamics simulations of full-length N dimers in the presence of RNA of varying lengths. A- Representative frames, obtained from CG simulation, of mean radius of gyration for N protein dimer in complex with a single strand RNA molecule with 10, 30 or 60 nucleotides. N protein monomers are colored in red and green whereas the RNA is colored in yellow. B- Density plot of the gyration radius distribution (in nm) calculated from five independent CG simulations of the N protein dimers in the presence of single-strand RNA of 10 to 70 nucleotides (nt) long. The median of distribution is indicated by a vertical line. C – Density plot of the frequency of contacts between N protein and any RNA nucleotide during the CG simulation. For instance, a frequency of 100% for N protein residue 260 means that this residue, from any of the N protein monomers, makes contact with at least one nucleotide of the RNA segment in all the analyzed frames of the CG simulations. The distance cut-off used to consider a protein-RNA contact in CG model was 10 Å between the alpha carbon of the protein residues and any atom of the RNA.

To better correlate the CG simulations with the negative stain images, we built a low-resolution 3D density map from the toroid-like particles (Figure 5). The comparison between the class averages used to build the 3D map and reprojections generated from the map is presented in Figure S4. By fitting the atomic model derived from the CG simulations performed with the 60 nt-long RNA into the 27 Å resolution density map (Figure S5A), we found that the NTDs are oriented side-by-side facing the CTD dimer (Figure 5B). All domains, including the flexible regions, contribute to shield the RNA segment. Interestingly, according to this model, there are no significant interface contacts between the CTD dimer and the NTDs (Figure 5, B and C). As already mentioned, most class-1 particles display a weak density at the center, which indicates the existence of empty space between the globular domains (Figure 5A) and corroborates the idea that these domains do not fully interact with each other [47]. It is noticeable that, in the proposed model, the interaction of the RNA with the NTD resembles the binding mode of the NTD to a 10-mer RNA reported recently [23] (Figure 5C, upper inset). Likewise, amino acid residues implicated in RNA binding in the CTD [17,22] are also involved in RNA interaction in our model (Figure 5C, lower inset). In addition, the intrinsically disordered regions represented by the N-terminal end and SR linker also make contacts with the RNA (Figure 5B). Interestingly, according to the model, the arginine and lysine residues of the SR linker interact with RNA while the serine residues, putative sites for phosphorylation, remain solvent-exposed and thus accessible to protein kinases (Figure 5B).

**Figure 5.**
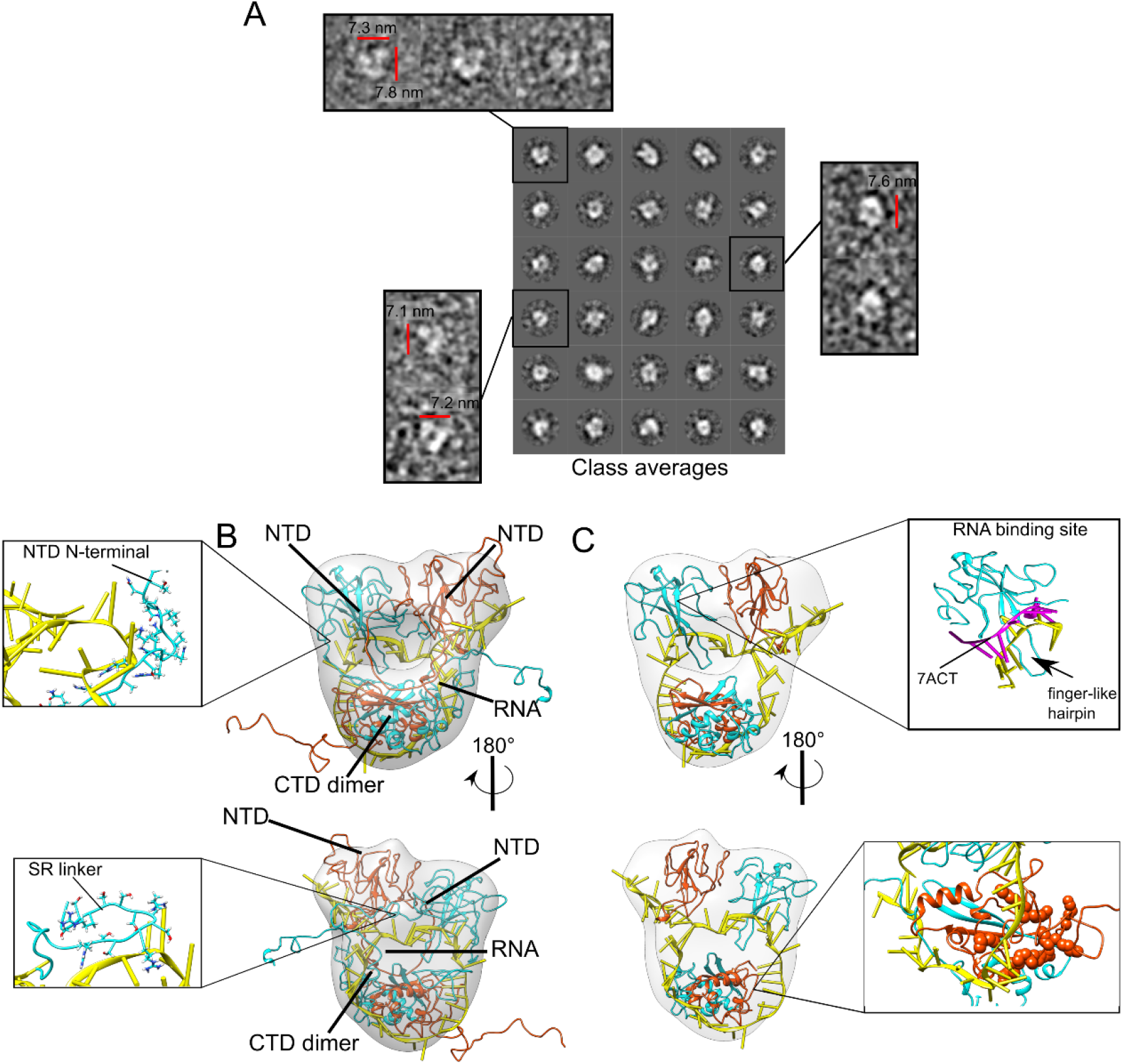
The full-length N dimer undergoes domain compaction in the presence of RNA. A- 2D classification of 24137 toroid-like particles picked with xmipp3. Thirty class averages used for the 3D model are shown in the main panel, whereas raw projections of particles representing the classes are presented in the insets. B- 3D density map reconstruction of the toroid-like particles with the flexible fitting of the N dimer atomic model derived from the CG simulations performed with the 60 nt-long RNA. Upper inset shows interaction between NTD N-terminal (residues 26 to 42 in sticks) and lower inset shows SR linker (residues 183 to 195 in sticks) contact with RNA. One N protein monomer is colored in orange and the other one is colored in cyan. RNA is colored in yellow. C- Same as B, but only presenting structured regions (NTD and CTD) of N dimer without its non-structured regions (NTD tail, SR-linker and CTD tail). The two NTD units are represented above the CTD dimer. Upper inset shows the superposition of the NTD from the structural model with the NMR NTD structure complexed with a 10-mer RNA in magenta (PDB code 7ACT [23]). Lower inset shows, as spheres, positively charged residues of the CTD region (248-280) that was previously demonstrated to be implicated in RNA binding [28,36].

Together, molecular simulations and 3D reconstructions from NSEM images suggest that full-length N undergoes domain compaction in the presence of RNA and offer an interpretation of how the NTD and CTD pack together in the protein dimer.

### The octameric model of full-length N bound to RNA

In addition to the toroid-like particles described above, we inspected the class-2 square-like particles, because these particles were also commonly found in the RNA-containing N preparations and presented a clear organization pattern with four rounded units connected by the edges (Figure 3B and Figure 6A).

**Figure 6.**
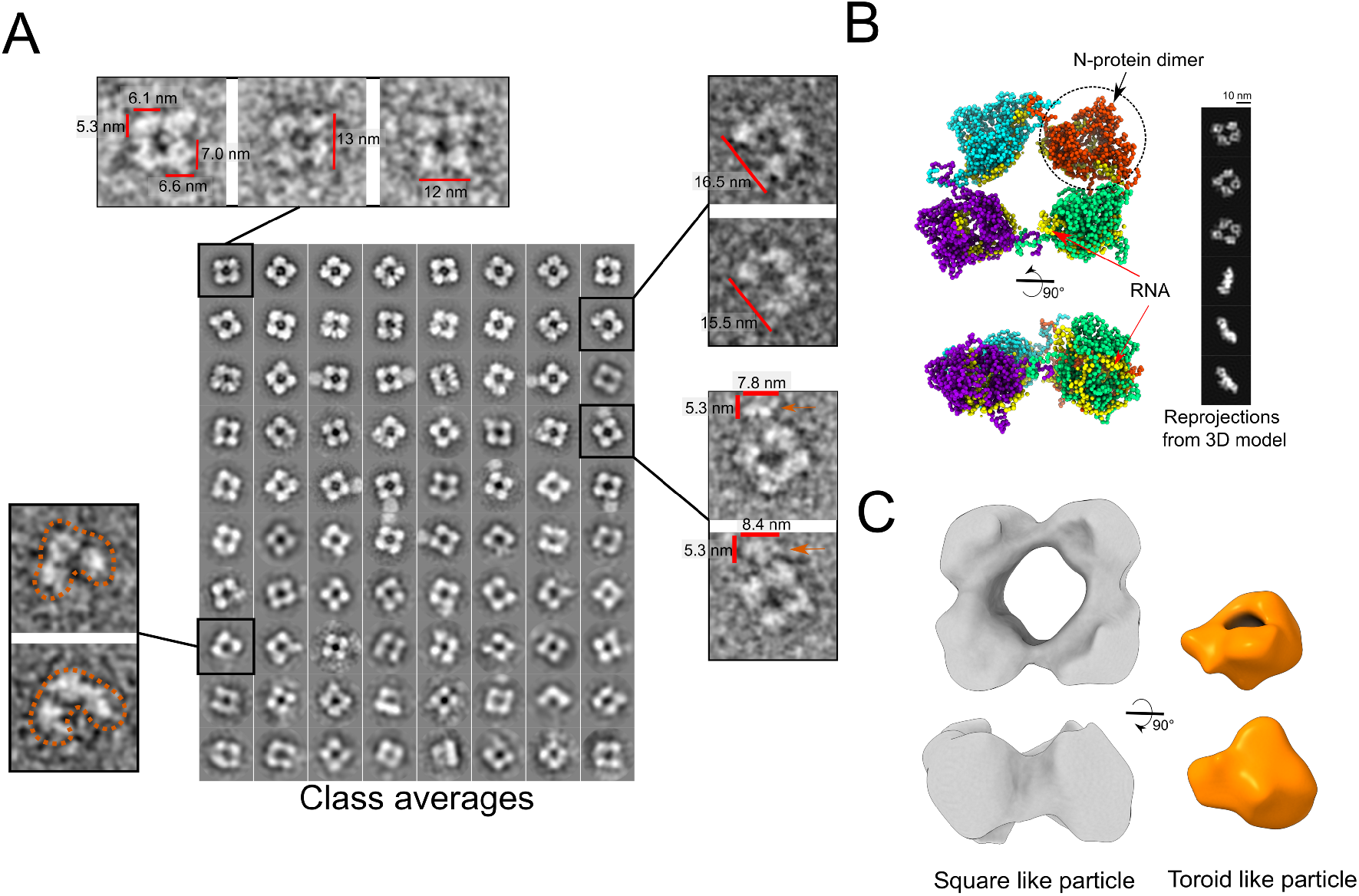
2D Classification and 3D density map reconstruction of the square-like particles. A- 2D classification of square-like particles picked with xmipp3. Class averages are sorted by the number of class members (number increase up and left). Raw projections of particles that compose the classes are shown on insets. B- Last frame of N protein octamer CG simulation in the presence of 60-nt RNA for each dimeric unit. Reprojections from the 3D structure are shown on the right. C- 3D-density map of the square-like particles in comparison to the map of toroid-like particles, both rendered at 3-sigma contour level.

To understand the structural organization of such particles in more detail, we collected ~25.000 class-2 particles and classified them into 150 subclasses (Figure 6A). These particles are ~13 nm wide and their rounded units located at the square vertices had dimensions varying from 5 to 7 nm. The size of these rounded units is consistent with the dimensions measured in the most populated class averages, the toroid-like particles (Figure 6A). The dynamic nature of the full-length N protein is highlighted by structural variations even within the same class (Figure 6A).

The 2D class average analysis also revealed particles showing a gap between two adjacent rounded units, like a U-shaped particle (Figure 6A, right inset). Of notice, several square-like particles showed a blurred appendix of similar dimensions as the toroid-like particles (Figure 6A, right inset). Other well-defined patterns comprise particles that seem to be composed of only three rounded units (Figure 6A, left inset). These findings suggest that the square-like particles are formed by independent N protein units, probably dimers compatible with toroid-like particles. Moreover, the NSEM images illustrate the pleomorphic nature of the N protein oligomers. As non-identical particles are an obstacle to finely uncover the N protein structure through single particle averaging protocols, we used the CG analysis to generate 2D template reprojections. These simulated reprojections led a new NSEM image particle picking and rationalized a 3D reconstruction of the squared particles, providing a possible N protein structural organization of high order oligomerization.

Thus, considering that each of the globular corners of the square-like particles has dimensions of 6 nm across and are formed by four independent N protein units, we modeled four copies (octamer) of the most packaged N-protein dimer from the 60-nt-long RNA simulation by placing each copy at the vertices of a virtual square. Then, in CG simulations, we connected each N protein dimeric unit through their C-terminal tails, based on the role of these tails in N protein oligomerization [25–27,48] (see next session and methods for details), and allowed approximation and accommodation of the oligomeric system (Figure 6B).

From the CG octameric structure, we built a simulated density map at 20 Å resolution, close to the resolution of the negative stain images, and generated reprojections from the 3D map at different orientations (Figure 6B). Remarkably, some reprojections are like the top view of the square-like particles and their dimensions. Further, reprojections corresponding to the side view of the octameric structure (Figure 6B) are consistent with the NSEM images of class-6 and class-7 particles (rod-like particles, Figure 3B). This finding was crucial because suggested that square and rod-like particles correspond to different orientations (top and side view, respectively) of the same particle in different orientations. Thus, by merging the original square-like particles with the set of rod-like particles, we built a low-resolution (26 Å) 3D density map for the square-like particles (Figure 5C and Figure S5). The 3D reprojections and original particle projections of both square and rod-like particles match appropriately, validating the reconstruction (Figures S6).

This map revealed four quasi-globular units connected through the edges in a planar configuration (Figure 6C). The volume of each globular unit resembles the volume of the toroid-like particles (Figure 6C), thus reinforcing the idea that each globular unit contains an N dimer. This structural organization is similar to the RNP particles purified from the MHV3 and HCV229E strains observed by negative staining [44]. Taken together, our data suggest that the square-like particles observed in NSEM images represent the top views of octamers of the N protein.

### van der Waals forces guiding the C-terminal tail self-interaction

The ability of the N protein to form dimers, tetramers, and higher-order oligomers in solution has been reported previously [17,22,27,36]. Nevertheless, how exactly N dimers interact with each other to form such higher-order oligomers is presently unknown. The N protein octameric organization proposed in the section above is quite complex and despite our density map being very informative, this map did not inform atomic details of the interaction involving N protein dimers.

*In vitro* studies suggest that the C-terminal tail residues 343-402 in SARS-Cov-1 and 365-419 in SARS-CoV-2 are required for N protein tetramerization [25–27,48]. Considering the high degree of identity shared by these regions between SARS-CoV-1 and SARS-CoV-2, a more restricted dimer-dimer interaction zone, comprising residues 365 to 402, can be reasoned. This region (365 to 402) is predicted to be unstructured in all coronavirus N proteins (Figure S7). Its adjacent C-terminal end residues, predicted to form an alpha-helix, are also conserved in most coronaviruses (Figure S7). Thus, in order to identify potential protein-protein contacts involved in oligomerization, we performed all-atom molecular dynamics simulations of the C-terminal tail.

One μs simulations starting with separated C-terminal tail monomers, *a* and *b*, revealed that the two monomers adopt a similar folding, retaining two alpha helices named α1 (375-382) and α2 (400-419) (Figure S8), consistent with the secondary structure prediction (Figure S7). The simulations also revealed intrachain contacts in each C-terminal tail monomers (Figure 7, A and B). *Monomer a* showed persistent van der Waals interactions between residues 407-415 (helix α2) and residues 377-385 (helix α1) and its neighbor coil residues L382 and R385 (Figure 7, A and D). The contacts involving helix α2 (N406, L407, S410, M411, S413, A414) with the coiled region of *Monomer b* (mainly residues P383, V392, L395) were also observed (Figure 7, B and E). C-terminal tail dimer formation was observed in all the simulations and the interface was asymmetrically formed by helix α2 from one of the monomers (Figure 7F). Overall, the results suggested that van der Waals interactions involving mainly the hydrophobic sequence LLPAA (394 to 398) (Figure S9) were the major forces driving the C-terminal tail dimerization.

**Figure 7.**
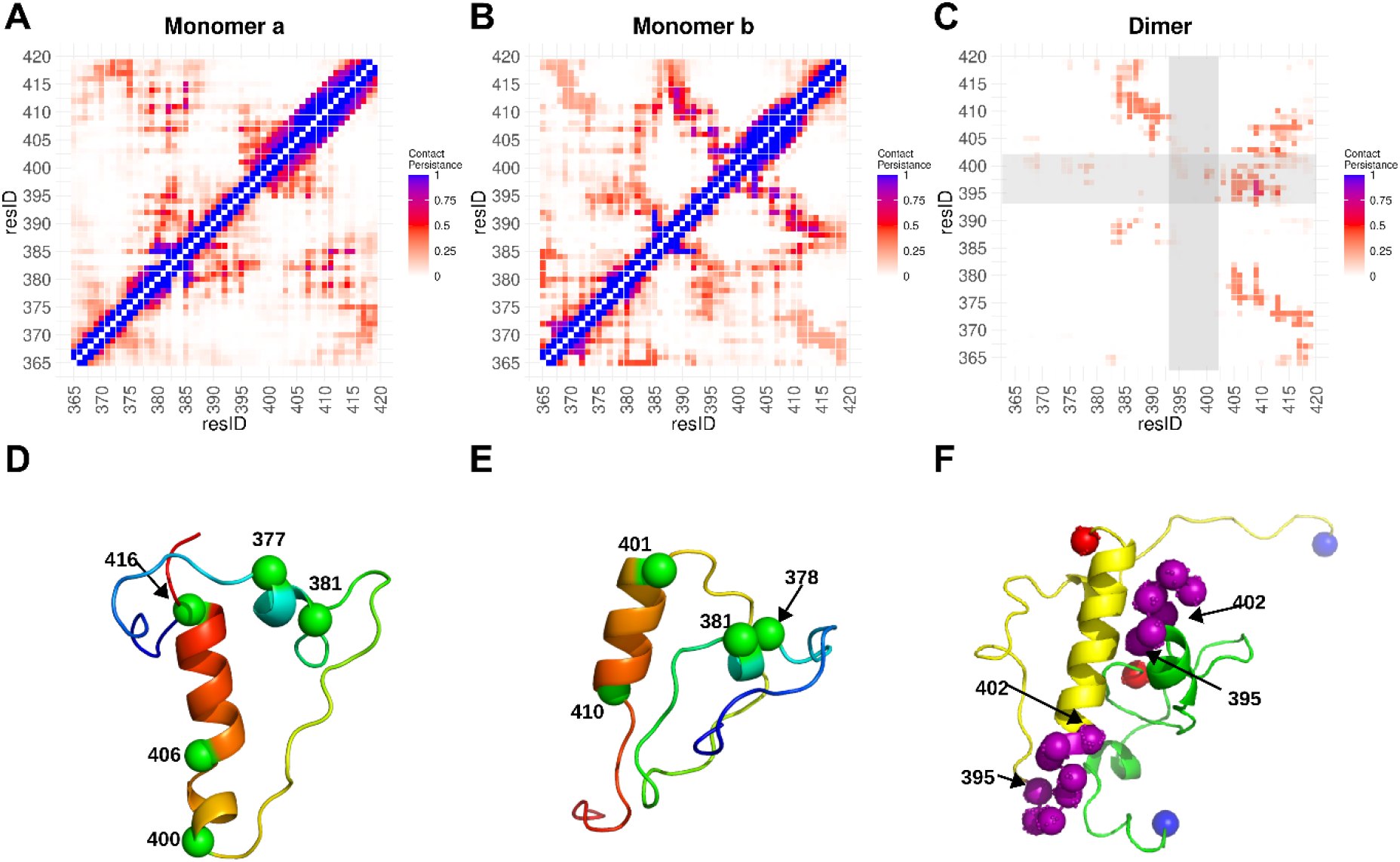
All-atom molecular dynamics simulation of the N protein C-terminal tail. A- and B- Contact maps for each individual monomer (a or b) or C-for C-terminal tail dimer. D-, E- and F- Representative structures of contact maps. The monomer contact maps (a or b) is an average of the whole trajectory of five MDs, considering only the last 50 ns of each trajectory for making the dimer contact map. The scale is defined as the contact persistence from 0 (none) to 1 (along all the simulation). D- and E- Structure of each monomer colored from N-terminal in red to C-terminal in blue, in green VDW some Cα to reference residues number. F- A representative structure of the dimer. Each monomer is in new cartoon representation, one in yellow and the another in green. In purple, the hydrogen atoms from the residues 395 to 402 suggested to participate in the dimer interface (grey transparent regions in C, see [27]). Spheres in blue and in red indicate the n- and c-terminal of the C-terminal tail used in the simulation.

## Discussion

The underlying mechanism of how the N protein associates with the RNA to form the RNP particles remains unknown. Likewise, the molecular basis of the N protein self-association is also poorly understood. The understanding of these processes, which are fundamental for the virus life cycle, has been hampered by the dynamic structural nature of the N protein. Thus, despite the wealth of structural data for the NTD and CTD regions, no structural models are available for the full-length N protein. Here, by combining NSEM with biochemical/biophysical and *in silico* approaches, we propose structural models and their dynamics to explain the manner of the dimer of the full-length N protein self-organizes both in the absence or presence of RNA. We also proposed a model of full-length N bound to RNA in an octameric organization that is reminiscent of RNP particles isolated from other coronaviruses [40,42].

The negative stain images of full-length untreated N bound to bacterial RNA provided an informative panel of N protein structural organization in the presence of large variety of nucleic acids. This is quite relevant because the structural organization of SARS-CoV-2 N protein is still poorly understood and only few studies have addressed this issue. The negative stain images showed a wide range of particles of different sizes and shapes, including large helical-like structures, which reflected the pleomorphic nature of this protein. Although particles of N protein incubated with polyA15 has a globular shape consistent with class 1 particles, thought to contain *E. coli* RNA, those derived from samples incubated with polyA50 showed irregular shape and were distinct from the well-organized class 2 particles, which are also believed to carry bacterial RNA. This reflected the dynamics of the N protein upon RNA binding and suggests that sequence-specific RNAs from *E. coli* can drive the packing of the N protein more effectively than the polyA molecules tested.

Despite N protein being typically described to form phase-separation [28,29], experimental conditions seem to be critical to induce such process. Here, the N protein preparations, even in the presence of RNA, did not present any evidence of phase-separation. The absence of phase-separation can be consequence of N protein concentration, which here is below the concentration demonstrated to induce phase-separation. Likewise, the RNA lengths used here (polyA15 or polyA50) are shorter than the one used in experiments to evaluate phase-separation [29].

Previous electron microscopy studies have suggested that the RNPs isolated from MHV and SARS-Cov-1 virions display a helical structure [41,42]. Furthermore, two studies using cryo-electron tomography have proposed that native SARS-CoV-2 RNPs are highly heterogeneous and densely packed but locally ordered in the virus, with neighboring RNP units with dimensions about 14 nm organized in a ‘‘beads on a string’’ fashion [43,49]. Moreover, although further studies are still required to confirm a helical organization of the SARS-CoV-2 RNP, the coiled structures of the MHV RNPs of about 11 nm in diameter with a 4 nm space [41] are quite compatible with the dimensions of the square-like (class 2) particles reported here. To our knowledge, these square-like particles have not been described for SARS-CoV-2 or any other coronaviruses N protein, although they are remarkably similar to electron microscopy images of RNPs isolated from MHV3 and HCV-229E [44]. We thus propose that the N protein octamers with the shape of the square-like particles described here could represent building blocks of the SARS-CoV-2 RNP structure, as suggested for the MHV RNP [42].

To form such higher-order structures with the RNA, the N protein is thought to also depend on protein-protein interfaces. Accordingly, the C-terminal tail of the coronavirus N proteins has been implicated in protein tetramerization [25–27,48]. Recently, C-terminal tails were reported to be crucial for the interaction between N protein dimers, and their deletion was sufficient to disrupt phase-separation [50]. Here, using molecular dynamics, we investigated how the C-terminal tail could play a role in the N protein oligomerization. We found that the C-terminal tail adopts a folded structure maintained by van der Waals intrachain contacts involving two helices. Moreover, the hydrophobic segment LLPAA appears to play an important role in tail-tail interaction, which agrees with HDX values for this segment reported previously (Figure 7F) [27]. The hydrophobic character of the C-terminal tail is thus thought to drive the formation of higher order oligomers (tetramer and octamers) of the N protein. A similar mechanism was found in Tula hantavirus N protein, in which the hydrophobic IILLF segment located at the C-terminal tail of its N protein was essential for the protein oligomerization [51]. The relevance of C-terminal tail as a target to affect N protein oligomerization was investigated in the related coronavirus HCoV‐229E, where a peptide derived from the tail was demonstrated to reduce the ability of N protein to form high-order oligomers [26]. In this sense, peptidomimetic molecules could thus be designed and used to disrupt this tail-tail interface to prevent N protein oligomerization.

The dynamic nature of N as an RNA-binding protein was also revealed by CG simulation models, which corroborated the NSEM images and radius of gyration of the protein in the absence and presence of RNA [19,36]. According to these models, in the absence of RNA, the N protein oligomerizes into dimers where the NTDs move freely relative to the CTD dimer, as previously suggested [36]. Conversely, in the presence of RNA, the protein undergoes a compaction that brings the NTDs closer to the CTD dimer. This compaction was confirmed by DLS measurements of the protein with polyA15. Despite the proximity of the NTD and CTD upon RNA binding, we did not observe a direct contact between these domains, as suggested by the weak density at the center of the 3D density map of the toroid-like particles, which resemble MHV dimeric N protein particles [42]. The CG simulations suggested multiple RNA binding sites in N protein dimer and the RNA-binding mode in N protein structured domains (NTD and CTD) indicated by our CG simulations is also consistent with the RNA-binding mode determined experimentally for the NTD [23], and with the predicted RNA-binding surface of the CTD [17,22]. The CG models also suggested a role of the SR-linker in RNA binding, which is in line with previous findings [9,14,19]. For instance, the highest frequency of RNA contacts observed in SR-linker during CG simulations occurs in the region of N protein R189 residue, which was previously proposed to play a critical role in RNA binding [29]. Also, mutations to positively charged amino acids in SR-linker close to this site, as the G204R mutation found in high-transmissibility SARS-CoV-2 lineage [31], could improve the ability of N protein binds to RNA. Considering that phosphorylation of the SR-linker plays a key role in the coronavirus life cycle [34], it is noteworthy that, in our CG models, the positively charged residues of the SR-linker point towards the RNA, while the serine residues are solvent-exposed and thus prone to be phosphorylated by protein kinases.

In addition, the N protein dimer models provided here offer further insights into how a single protein dimer interacts with a single RNA molecule of varying lengths without considering higher-order oligomerization. These models aimed to simulate how the N protein binds to the viral genome, where several protein dimers are thought to compete for a short genome segment. Our CG models suggested that a genome segment ranging from 40-60 nt would be sufficient to occupy all the RNA-binding sites and induce protein compaction. However, such RNA-binding sites could be simultaneously occupied by multiple short RNA segments, as suggested by the GC models and confirmed experimentally with polyA15 and polyT10 [37]. Likewise, longer RNA segments could drive protein oligomerization, as observed with polyA50 and polyT20 [37].

In conclusion, our results shed new light on the dynamics and higher-order oligomeric structure of the SARS-CoV-2 N protein and provide a framework for understanding the multifunctional and versatile role of this protein.

## Methods

### Cloning procedures

The SARS-CoV-2 RNA was isolated from virus particles with the QIAmp viral RNA mini kit (Qiagen - USA) and reversely transcribed to cDNA with the High-Capacity Reverse Transcription Kit (Thermo - USA). The N protein sequence (GenBank QIG56001.1) was amplified from cDNA samples using primers SC2-protN28182-F (5’-AGTCTTGTAGTGCGTTGTTCG-3’) and SC2-protN29566-R (5’-ATAGCCCATCTGCCTTGTGT-3’) and cloned into pGEM-T Easy (PROMEGA - USA), generating plasmid pGEM-SC2-N. The N sequence was reamplified from pGEM-SC2N with forward 5’-AACAAGCTAGCATGTCTGATAATGGACCCCAAAATCAG-3’ and reverse 5’-GGTCTGCGGCCGCTTAGGCCTGAGTTGAGTCAGCACTGCT-3’ primers and subcloned into the *Nhe*I/*Not*I sites of a pET28a-TEV vector carrying a 6xHis-tag and TEV protease cleavage site at the N-terminus.

### Protein Expression and Purification

The N protein was expressed in *E. coli* BL21 (DE3) cells (Novagen -USA) and purified by metal-affinity and SEC. Cells were grown at 37°C under agitation (200 rpm) in LB medium containing kanamycin (50 mg/L) to an optical density (OD600nm) of 0.8. Recombinant protein expression was induced by the addition of 0.1 mM isopropyl-thio-β-d-galactopyranoside (IPTG) for 16 h at 25 °C. After centrifugation, cells were resuspended in 50 mM sodium phosphate, pH 7.6, 300 mM NaCl, 10% v/v glycerol, 1 mM phenylmethylsulfonyl fluoride and incubated on ice with lysozyme (0.1 mg/mL) for 30 min. Bacterial cells were disrupted by sonication and the soluble fraction was loaded on a 5 mL HiTrap Chelating HP column (GE Healthcare - USA) previously equilibrated with same buffer. Proteins were eluted using a linear gradient (20 to 500 mM) of imidazole at a flow rate of 1 mL/min. Eluted fractions containing the N protein were pooled, concentrated and loaded on a HiLoad 16/60 Superdex 200 column (GE Healthcare), previously equilibrated with 10 mM Tris, pH 8.0, 100 mM NaCl, at a flow rate of 0.5 mL/min. These samples, still carrying bacterial RNA, were named as untreated N protein samples in the text.

To produce the N protein without nucleic acid contaminants we treated the purified N protein with RNAse A or we purified N protein in the presence of Urea and high salt concentration. To treat N protein with RNAse A, purified N protein was incubated with RNAse A (Qiagen) with 1:15 (RNAse A: protein) proportion for 1 h at room temperature. After incubation, the samples were loaded on a 16/60 Superdex 200 column equilibrated with 50 mM sodium phosphate, pH 7.6 and 500 mM NaCl. Eluted fractions containing the N protein were pooled and concentrated.

To produce urea-treated N protein samples, *E. coli* cells were lysed in buffer A (50 mM sodium phosphate, pH 7.6, 500 mM NaCl, 20 mM Imidazole, 6 M Urea, 10% Glycerol). The suspension was sonicated and centrifuged at 18,000 × g for 45 min at 4 °C. The supernatant was applied on a HiTrap Chelating HP column equilibrated with the same buffer. Proteins were eluted with a linear imidazole gradient using buffer B (50 mM Sodium Phosphate, pH 7.6, 500 mM NaCl, 300 mM Imidazole, 3 M Urea, Glycerol 10%). Protein fractions were mixed and dialyzed against buffer C (50 mM sodium phosphate, pH 7.6, 500 mM NaCl, 10% Glycerol). Protein samples were centrifuged at 14000 × g for 10 min at 4 °C and subjected to molecular exclusion chromatography on a Superdex 200 16/60 column, equilibrated in buffer C, under a flow rate of 0.7 mL/min. Protein purity was analyzed by SDS-PAGE and protein concentration was determined by absorbance at 280 nm using the molar extinction coefficient calculated from the amino acid composition. Protein samples were concentrated and stored at – 80 °C. In experiments involving polyA15 or polyA50, except for SEC-MALS analysis, urea-treated samples were diluted 5x (DLS) or 20x (NSEM) in 50 mM sodium phosphate, pH 7.6 buffer (without NaCl) prior to RNA incubation.

### Circular dichroism analysis

Protein samples at 6 to 30 μM final concentration were diluted in 50 mM sodium phosphate buffer, pH 7.6, and analyzed by FAR-UV circular dichroism. All measurements were recorded on a Jasco J-810 Spectropolarimeter at 10°C, in the range of 197-260 nm. After buffer signal subtraction, CD signals were normalized to residual molar ellipticity using the equation *θ* = (*mdeg*.100.*MW*)/ (*mg/mL.l.NR*), where mdeg = CD signal, MW = protein molecular weight, mg/mL = protein concentration in mg/mL, l = optical path in centimeters and NR = protein residues number.

### Dinamic Light Scatering

N protein samples (~ 20 μM) in 50 mM sodium phosphate, pH 7.6, 100 mM NaCl, 2% Glycerol, were submitted to dynamic light scattering analysis in the ZetaSizer NanoZS90 (Malvern) equipment. All the measurements were acquired following the equipment automatic setup, and after sample equilibrium at 10 - 18 °C. All buffer parameters corrections were performed by Zetasizer Nano software. Obtained data are shown as the average of at least three measurements. The statistical approach used to compare the conditions was performed in R with DTK R package that applies the Dunnett-Tukey-Kramer pairwise multiple comparison test adjusted for unequal variances and unequal sample sizes. The P value was obtained from the confident interval as described in [52].

### SEC-MALS analysis

N protein samples (~ 20 μM) were loaded onto a Superdex 200 10/30 column, equilibrated with 50 mM sodium phosphate, pH 7.6, 500 mM NaCl, under a flow rate of 0.4 mL/min. To inspect the RNA influence on the N protein oligomeric state, samples of N protein in 5-fold excess of RNA polyA50 or polyA15 were also analyzed following the same protocol. For this, the mixture (protein-RNA) was incubated, on ice, for 1h, before the injection. SEC-MALS analyses were performed on a Viscotek OmniSEC (Malvern, UK) equipped with a SEC module coupled to a two-angle laser light scattering detectors, a refractometer and a viscometer. The OmniSEC software was used to acquire and evaluate the data. All the graphs were obtained using OriginPro 2021 software.

### Negative staining and imaging

To collect negative stain images of the untreated N protein, 3 μL of purified N (4 μM) in 10 mM TRIS buffer, pH 8.0, 100 mM NaCl were applied onto glow-discharged (15 mA, negative charge for 25 s) 400-mesh copper grids covered with a thin layer of continuous carbon film (01824 - Ted Pella, USA). After 1 min, the excess liquid was drained using a filter paper. The grids were stained twice with 3 μL uranyl acetate solution (2%) for 30 s. The excess solution was drained and the grids were allowed to dry at room temperature. Automated data collection was performed using a 200 kV Talos Arctica G2 transmission electron microscope (Thermo Fisher Scientific). Data set of 18148 micrographs were automatically acquired with EPU software and recorded on a Ceta 16M detector. The pixel size and defocus were 1.96 Å and −1.5 μm, respectively. The exposure time was 1 s in an accumulated dose of ~23e-/Å^2.^

To collect negative stain images of N protein treated with urea in the absence or the presence of polyA15 or polya50 we used the same grid preparation protocol described above. Urea-treated samples in 50 mM sodium phosphate, pH 7.6, 500 mM NaCl, 10% Glycerol were diluted 20x in the same buffer but without NaCl before incubation for 2 h at room temperature with RNA. The protein final concentration was 0.5 and we used 1:10 protein-RNA molar ratio in N protein samples with RNA. Screening data was performed using a 120kV JEM-1400Plus transmission electron microscope (JEOL, Japan), equipped with OneView 16-Megapixel Camera (Gatan, USA), magnification of 80k and pixel size of 1.39-1.89Å.

### Image processing

A total of 17370 collected negative stained micrographs (4096 × 4096) from purified N protein (untreated-N) preparations were preprocessed using Imagic-4d software system [53]. Raw micrographs were firstly submitted to *a posteriori* camera correction and then to a contrast transfer function (CTF) correction. For CTF correction, the amplitude spectrum of each image was determined and submitted to eigenvector analysis and automatic unsupervised classification into 2000 classes. The resulting class averages were used to determine the CTF parameters, which were passed to the individual images. Then, the CTF correction was applied by phase-flipping each image. A total of 15474 CTF-corrected micrographs were selected based on the quality of CTF estimation and defocus for further processing. All image processing programs, except Imagic, were run in the Scipion framework [54].

Micrographs were resized in Fourier space to 1024 × 1024 dimensions and submitted to particle picking using the Xmipp3 package [55]. The picking was performed in original 4096 × 4096 size, by an initial manual picking of ~550 particles from 40 micrographs, and proceeded by automatic picking, resulting in 178,021 particles in a 200 × 200 box size. The particles were subjected to a round of 2D classification in Relion to separate toroid, square and rod-like particles groups. A total of 24137 toroid-like particles (class 1), 33550 square-like particles (class 2) and 12552 rod-like particles (classes 6 and 7) were used for further processing.

Given the structural pleomorphism of purified N protein, we cautiously evaluate the use of classical single-particle protocols. Toroid-like particles were classified into 300 class averages using Relion [56] and an initial 3D model (C1 symmetry) was built using 30 class averages in Eman2 [57]. Class averages placed at a 112 × 112 box with a circular soft mask (0.52 radius and 0.05 drop-off) were band-pass-filtered (0.001 low pass and 0.1 high pass). Then, the initial 3D map was refined in Imagic using an iterative process of angular reconstitution and class average rotation and translation alignment to 3D reprojections, achieving 3D resolution convergence.

In the case of square and rod-like particles, these particles were cautiously merged into a combined particles dataset being informed by the 3D structural models obtained from CG molecular dynamics simulations of N protein octamer. From these particles, an *ab initio* reference-free initial volume was generated in Relion, followed by two independent rounds of 3D classification. We calculated the consensus [58] of both 3D classifications (resulting in a total of 9 3D classes) and selected a single stable 3D class corresponding to 17,803 particles (38.6% of the particle set). This 3D class was further auto-refined following the 0.143 FSC target in Relion, obtaining self-consistent convergence in 13 iterations. The 3D density unmasked map resolution was accessed by the Fourier Shell Correlation (FSC), using ½-bit threshold (Figure S4).

### CG molecular dynamics simulations

CG molecular dynamics simulations were performed using CafeMol 3.1 software [59]. An initial N protein dimer all-atom model was built in YASARA software [60] using crystallographic structures of NTD monomer (PDB ID: 6VYO) and CTD dimer (PDB ID: 6WJI). For this, NTD monomers were placed at a distance that allowed a fully extended NTD-CTD linker conformation. Intrinsically disorders domains (N-terminal tail, NTD-CTD linker and C-terminal tail) were modeled using YASARA. The modelling was based on the N protein sequence from GenBank QIG56001.1, the same used in experimental assays. For simulations in the presence RNA, an initial single strand RNA all-atom model was modeled without tertiary structure or base pairing in YASARA. Nucleotide sequence of all RNA of the same size used in simulations were the same and consist of a scrambled sequence, since the CG model used does not compute base specific interactions.

Five independent simulations with different random seeds were conducted for each dimeric condition (without RNA or in the presence of a single RNA segment of 10 to 70 nucleotides) by Langevin simulations. For each independent simulation, the RNA was initially placed at different aleatory positions in relation to the protein. The temperature was set to 300 K. Local protein-protein and RNA-RNA interactions were modeled using AICG2+ and GO potentials[59], respectively. For non-specific interactions between N protein monomers, and between N protein monomers and RNA, excluded volume and electrostatic interactions were considered. Electrostatic forces were computed using Debye–Hückel equation with a cutoff of 20 Å and ion strength of 0.1 M. To positively charged residues +1e was assign whereas for negatively charged –1e. Local GO potential was changed to Flexible Local Potential in intrinsically disorder regions: NTD tail, SR-linker and CTD-tail. Each simulation was run for 1.5×10^7^ steps, with a time length of 0.2 for each step in CafeMol scale. For dimers simulations, only the last 5×10^6^ were used in MD analysis.

For N protein octamer simulation, we built an initial model using four N protein dimer copies from the last all-atom fitting model. Each dimer was placed 200 Å from the center of mass of the neighbor dimer. In CafeMol simulations, a harmonic spring (force coefficient of 0.5 and distance constraint of 5) was used to bring together C-terminal tails from neighbor dimers. The contacts involving each dimer was modeled using GO potential to maintain native structure of both protein and RNA. In addition to the harmonic spring, we used excluded volume and electrostatic interactions to model the contacts among the four dimeric units. The octamer simulation was run for 5×10^6^ steps, which was sufficient to stabilize the RMSD calculated from CA atoms. The last frame from the simulation was further analyzed.

### Tools for comparing CG simulations with biophysical or electron microscopy data

For comparing CG simulation radius of gyration with biophysical data, we estimated the radius of gyration of the simulated systems with Bio3D package using an in-house R script [61].

For comparing 3D atomic structures or CG models with negative stained images, we reprojected the structures or the models into 2D simulated images. Then, we estimated the area of these simulated reprojections as well as the area of the particles from negative stained images. For the full-length N protein dimer, we used 3 structures obtained from CG simulations: structures with the minimum, mean and maximum radius of gyration along the simulation. Prior to reproject, CG structures were converted to all-atom structures. For NTD monomer and CTD dimer structured domains, we used crystallographic structures of these domains obtained from Protein Data Bank (PDB ID: 6M3M [24] and 6WZO [27], respectively). All the structures, obtained from CG simulations or PDB, were used to build 20 Å resolution simulated density maps with Chimera [62] using *molmap* function. Then, the 3D density maps were reprojected in different orientations (rotational angle 0 to 360 degrees with step of 20 degrees and tilt angle 0 to 180 degrees with step of 20 degrees) with Xmipp3 *create gallery* function in Scipion workflow. A total of 100 2D reprojected images were used for each structure and, before any comparison, images were resized to match the pixel size values.

To estimate the area of simulated reprojections of 3D structures or the area of the particles from the negative stained images. we used ImageJ software [63]. In ImageJ software, we first adjusted a threshold to indicate the region of the image corresponding to N protein, as showed in Figure S2B. In the case of the full-length N protein dimer from CG simulations, its simulated reprojection can be seen as three high contrast regions, depending on its orientation and conformation (Figure S2B). These regions correspond to the N protein structured domains (NTD and CTD), and in this case, these regions were computed separately in the area estimation process.

### 3D Atomic model fitting

From the previous independent simulation of the CG model of N protein dimer in complex with a 60-nt-long RNA were selected 100 structures with the lowest radius of gyration using only the structures domains (NTDs and CTDs). The correlated structures were removed by clustering. The structures of reduced set (~40 structures) were converted to all-atom structure model. The protein was reconstructed using PULCHRA[64] and the RNA was converted into all-atom model using DNAbackmap tool from CafeMol.

The structures were rigid-body docked on the density map by employing *colores* program from Situs package[65]. Three different docking calculations were performed by selecting three different structural segments of the system: 1) NTDs, CTDs and RNA heavy atoms; 2) NTDs and CTDs heavy atoms, and 3) all heavy. The docked structures at each group were ranked by its cross-correlation coefficient (CCC). The selected structure for the model fitting was that with the lowest CCC and with close spatial coordinates in all the docking calculation groups. A compatible resolution with the density map (25 Å) was used for the conversion from PDB to volumetric map.

The VMD AutoPSF plugin was used to generate a topology for MD simulations and MDFF simulations. The complete all-atom model (protein-RNA) was solvated in a TIP3P water box of 134×191×163 Å^3^ using Solvate plugin from VMD. Na^+^ and Cl^−^ ions were added to neutralized and to adjust the ionic strength to 150 mM using the VMD plugin autoionize[66]. The complete system involved 396,013 atoms. The molecular interactions were described by CHARMM36 force field with CMAP corrections[67].

All simulations were performed with NAMD 2.13[68]. The system was equilibrated using the following protocol: 1) 10,000 minimization steps of water molecules and ions by restraining the protein and RNA; 2) 10,000 minimization steps of the complete system; 3) 200 ps of NVT MD fixing positions of the protein and the RNA and 4) 200 ps of NPT MD fixing positions of the protein and the RNA. The temperature was maintained in 300K using a Langevin thermostat with damping coefficient of 1 ps^−1^ and coupling only heavy atoms. In the NPT simulations, the pressure was maintained constant in 1 atm using Nose-Hoover barostat with a piston decay of 200 fs and a piston period of 400 fs. Particle Mesh Ewald (PME) method were applied for describing long-range electrostatic interactions[69]. A non-bonded cut-off of 10 Å was applied to calculate electrostatic and van der Waals potential (vdW). A shifting function was applied to electrostatic interactions for avoid truncation of the potential function. vdW interactions were smoothed by applying a switch function at distance 9 Å. The time step was 2 fs using SHAKE for all covalent hydrogen bonds of the protein and nucleic molecules[70] and SETTLE algorithm for rigid water molecules[71].

MD Flexible Fitting of the protein-RNA into the density map were performed following a multi-step protocol[72]. The protocol involved three steps: 1) 5 ns MDFF simulation, applying a scaling factor used in the energy potential of the map (ξ = 0.3 kcal mol^−1^) on RNA and NTDs and CTDs. Also, the dihedral angles of α-helices and β-sheet of the NTDs and CTDs were constraint using harmonic restrains (kμ,protein= 400 kcal mol^−1^ rad^−2^). 2) 5 ns MDFF using similar conditions of the step 1, but including the intrinsically disordered regions (N- and C-terminal tails and SR-linker) in the potential energy function of the map. 3) a minimization of 10,000 steepest descent steps, just restraining backbone position of the protein and the RNA with a force constant of 10 kcal mol^−1^ Å^−2^. 4) a minimization of 10,000 steepest descent steps without any restraint. The time step was 1 fs. SETTLE algorithm was used for rigid water molecules[71]. Each step was done until reach convergence of the cross-correlation coefficient between model and map, and convergence of the RMSD calculated from protein and RNA backbone (Fig S10).

### All-atom molecular dynamics simulations

For C-terminal tail simulations, a folded linear protein was built with YASARA[60], from PSIPRED secondary structure prediction[73]. This structure shows one helix from residues 403 to 419 and was used as starting point for MD simulation of one C-terminal tail monomer. The MD simulation of the monomer was to check the folding along the trajectory without the influence of another monomer. The initial secondary structure did not change along the trajectory and no folding was observed. When the trajectory is aligned using the helix as reference, the coil region moves randomly. After a clustering, the average structure of the simulation was used as starting point for building a dimer system with the monomers separated by ~70 Å using Packmol program[74]. The large distance was set in order to decrease any possible bias on the starting configuration.

The dimer was immersed into an octahedral box with 56,098 TIP3P water box, 150 Na^+^ ions and 152 Cl^−^ ions. The dimer formation was study by running five replicates. However, this system was described by ~170,000 atoms, needing an important computational effort to perform a 1 μs MD simulation. Thus, five uncorrelated starting points of the dimer were picked from the dimer MD simulation. The criterion was to select those with a minimum distance of 30 Å between any atom of the monomers. All the selected dimers were solvated again in octahedral box with 45,556 TIP3P waters, 121 Na+ ions and 123 Cl-ions in order to keep the system composition and the ionic strength (150 mM). All the systems were described using Amber ff14SB force field[75] and the topologies were generated using *tLeap* program from AmberTool20[76].

All the MD simulations were equilibrated by the following steps: 1) minimization of the solvent by 2500 steepest descent steps followed by 2,500 conjugate gradient steps; 2) minimization of the whole system by 2,500 steepest descent steps followed by 2500 conjugate gradient steps; 3) the system was heated from o to 300 K in 200 ps under NVT conditions, restraining the protein atom positions; 4) the density was equilibrated under 500 ps under NPT conditions, restraining the protein atom positions. The production step of all the simulation was run for 1 μs under NPT conditions. For all the simulations, the temperature was set to 300 K using a Langevin thermostat with a collision frequency of 5 ps^−1^. In the case of the NPT simulations, the pressure was set to 1 atm using a Monte Carlo barostat with pressure relaxation time of 1 ps^−1^. The long-electrostatic interactions were calculated using Particle Mesh Ewald method[69] 9 Å. SHAKE algorithm was applied to all bonds, allowing a time-step of 2 fs. The simulations were run using GPU accelerated PMEMD program that is part of the AMBER18 package[77–79].

The contact map analysis was done using *nativecontacts* command using two masks, one of each monomer, including all atoms and a cut-off distance of 7 Å. The contact maps figures were done using Rstudio[80]. The secondary structure along trajectory was performed using *secstruct* command. The inter-monomers vdW and electrostatic interaction energy was calculated using *lie* command with a dielectric constant of 78. All the commands are part of Cpptraj program from AmberTools20[76]. The images of the C-terminal tail monomers and dimers were generated using pymol 2.3.0 (open-source build).

## Acknowledgments

We thank LNNano/CNPEM for the use of electron microscopy facility (TEM-26919, TEM-27882). This work is part of the Rede Virus MCTI taskforce on COVID-19 funded by FINEP (grant number 01.20.0003.00), Brazilian Ministry of Science, Technology and Innovation. The authors acknowledge financial support from the Fundação de Amparo à Pesquisa do Estado de São Paulo (FAPESP), project number 17/18139-6. We thank David A. Case from Rutgers University for the AMBER18’s license fee waiver.

## Author contributions

HVRF and GEJ performed all-atom and coarse-grained molecular dynamics simulations; GEJ performed molecular dynamics flexible fitting simulations; HVRF, GRS and SLG processed negative stain electron microscopy images; FAHB and CCT conducted biophysical experiments; ACB, AC, ASS, CCT and SLG prepared electron microscopy grids; ACB and AC collected electron microscopy images; FAHB, CCT and ASS expressed and purified proteins; MCB, REM, DBBT, KGF, ACMF, CEB and PSLO designed the experiments, analyzed the data and wrote the paper.

