## Supplementary Material for "Structural dynamics of SARS-CoV-2 nucleocapsid protein induced by RNA binding"

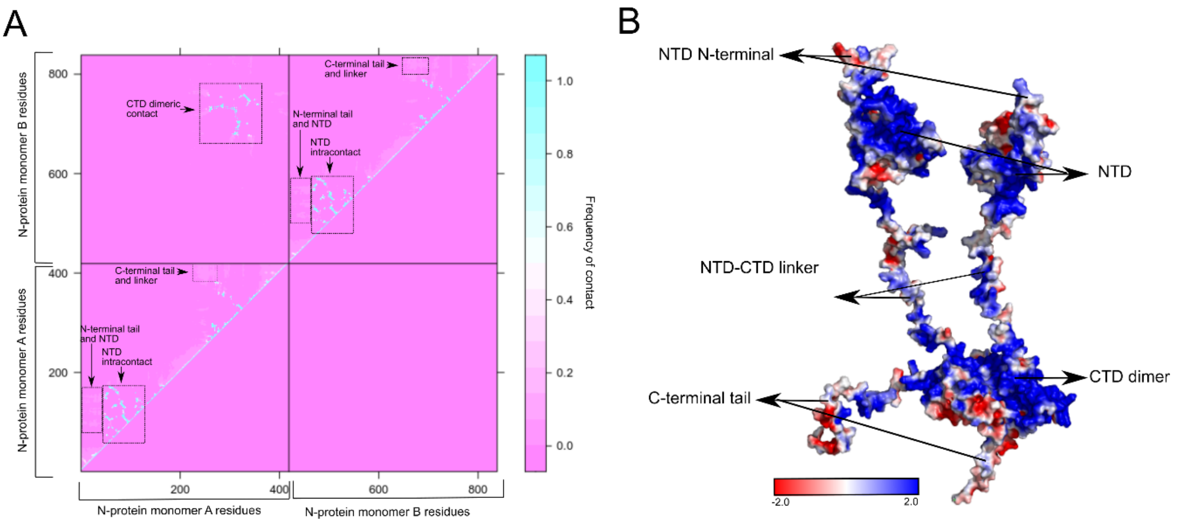


**Figure S1.** A- Contact map of the N protein dimer in RNA-free CG simulation. Contact map was built using Bio3D packaged. Five independent CG simulation runs were considered during frequency contact calculation and a cut-off distance between CG Cα beads was set to 10 Å. Relevant contact regions are indicated. B- Electrostatic potential surface of an extended conformation of the N protein dimer model.

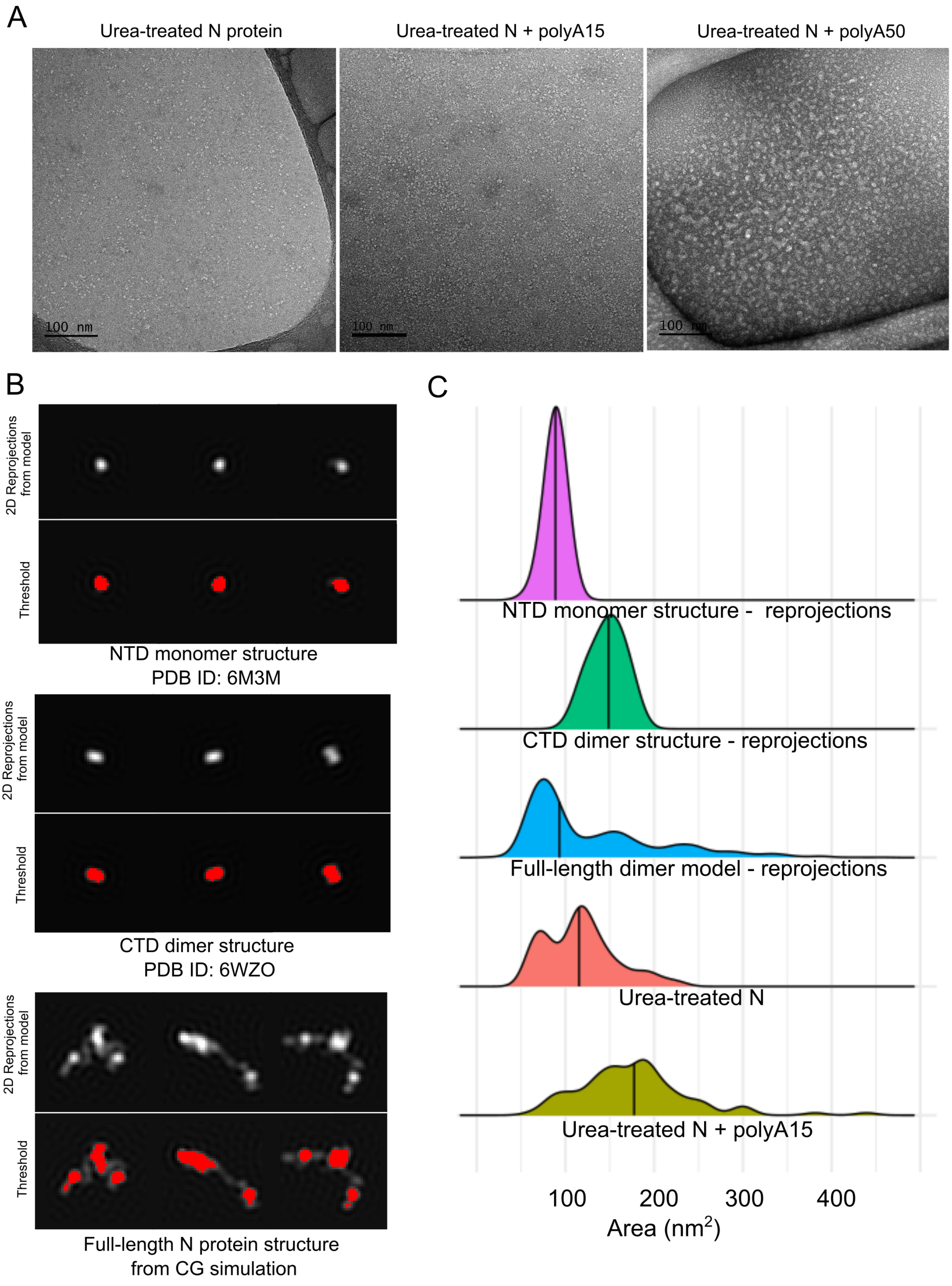


**Figure S2. Representative NSEM images of the N protein in the absence or presence of RNA and the area estimation of particles.** A- Representative micrographs for each condition: urea-treated N protein in absence of RNA (left panel), or in the presence of polyA15 (middle panel) or polyA50 (right panel) at a 1:10 protein-RNA ratio. Scale bars are indicated. B – Representative 2D simulated reprojections in three different orientations of NTD monomeric crystallographic structure obtained from PDB 6M3M (upper panel), CTD dimeric crystallographic structure obtained from PDB 6M3M (middle panel) and full-length

N protein structured obtained from CG simulations (bottom panel). Below each reprojections are presented the threshold applied to identify the high-density regions in images, which corresponds to the structured domains of N protein. In full-length N protein reprojections, under the threshold used we did not include the flexible regions of N protein that has low intensity value. C – Area distribution of 3D structures (NTD monomer and CTD dimer) or 3D model reprojections (full-length dimer) and class averages of particles collected from negative stain images of urea-treated N samples, same as in Figure 2B. Here, the area distribution of urea-treated N with polyA15 samples, were included in the comparison presented in Figure 2B of the manuscript. The area distribution of urea-treated N overlaps the area distribution of NTD and CTD reprojections, indicating that the particles observed in the negative stain images corresponds to these individual domains. The presence of polyA15 shifted the area distribution of urea-treated N, indicating that the particles have larger dimensions than without RNA, but still compatible with the larger dimensions observed in full-length dimer reprojections. This finding suggests that in presence of polyA15, the individual NTD and CTD domains got closer and the individual density regions that are separated, got together and formed a unique particle with larger dimensions in comparson to the individual domains. A 2D classification of the N protein + polyA50 particles could not be performed because these particles were not well delimitated.

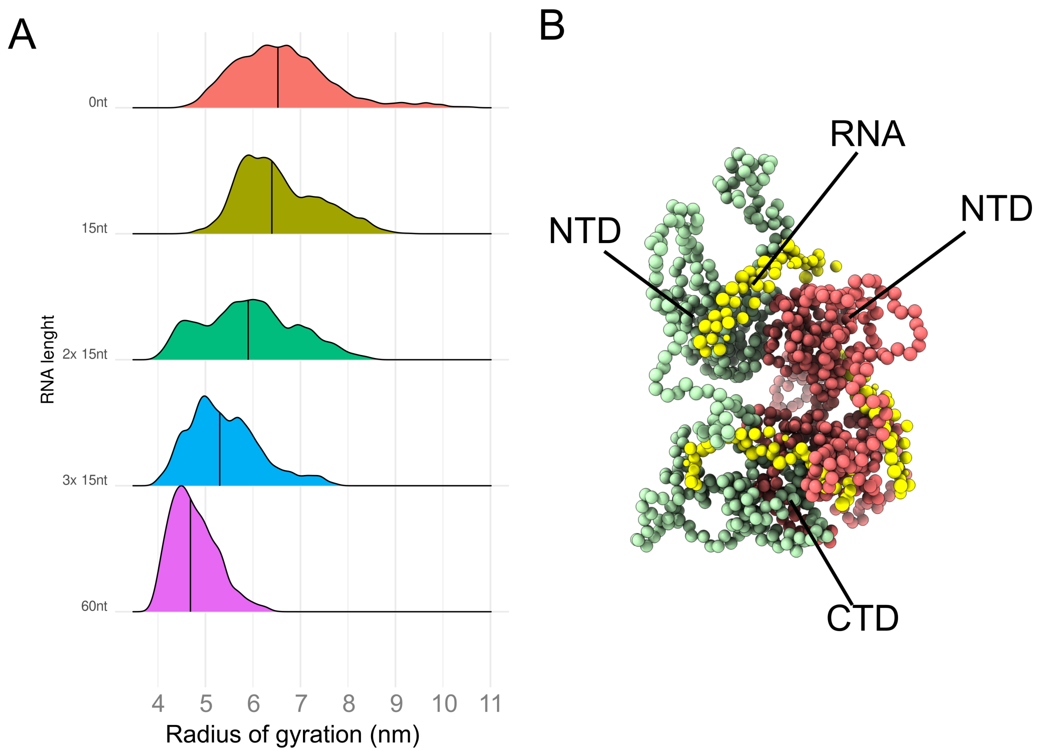


**Figure S3. CG dynamics simulations of full-length N dimers in the presence of 15-nt RNA molecules.** A- Density plot of the gyration radius distribution (in nm) calculated from five independent CG simulations of the N protein dimers in the presence of one, two or three single-strands 15-nt RNA. For comparison, we also presented distributions of conditions without RNA (0 nt) and in the presence of 60 nt long. B- Representative frame with minimum radius of gyration of full-length N dimer in complex with three 15-nt RNA molecules. N protein monomers are colored in red and green, whereas the RNA molecules are colored in yellow.

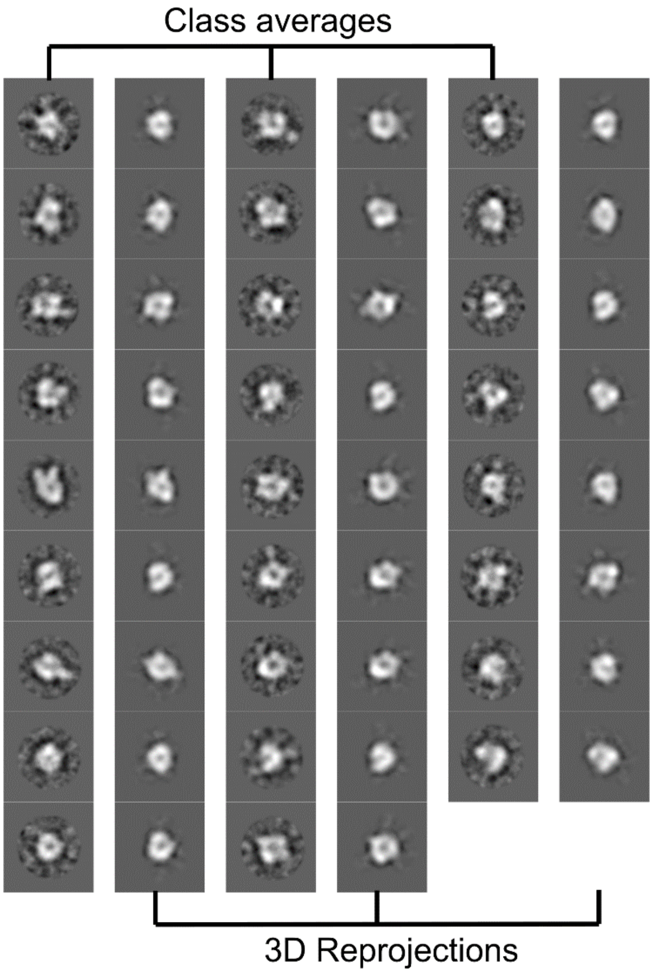


**Figure S4.** Comparison between toroid-like class averages (even columns) and their reprojections from the 3D volume obtained in EMAN2 (odd columns) to validate the 3D reconstruction.

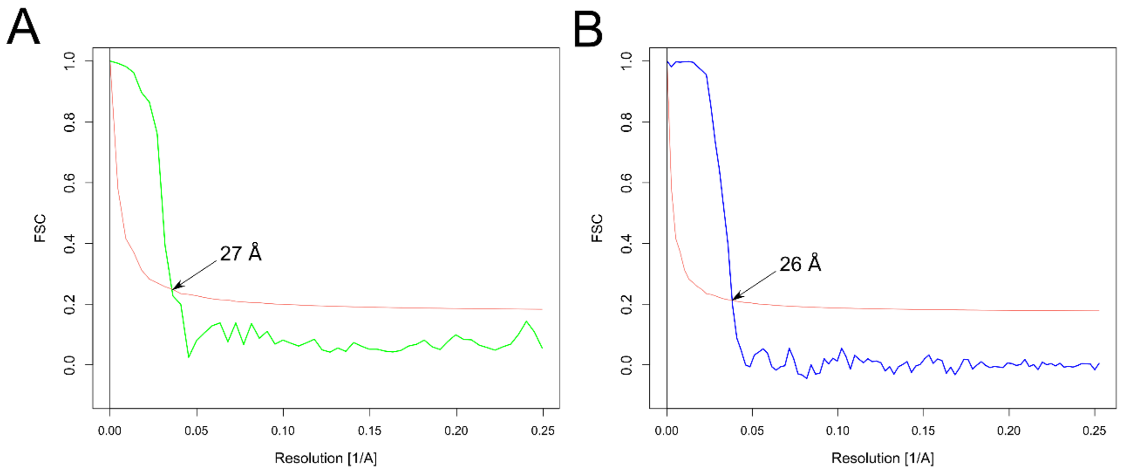


**Figure S5.** Fourier shell correlation profile of (A) toroid-like particles or (B) square-like particles. FSC curve calculated from two 3D volumes is shown in green or blue and the half-bit curve in red. The global resolution determined based on the half-bit criterion is indicated [1].

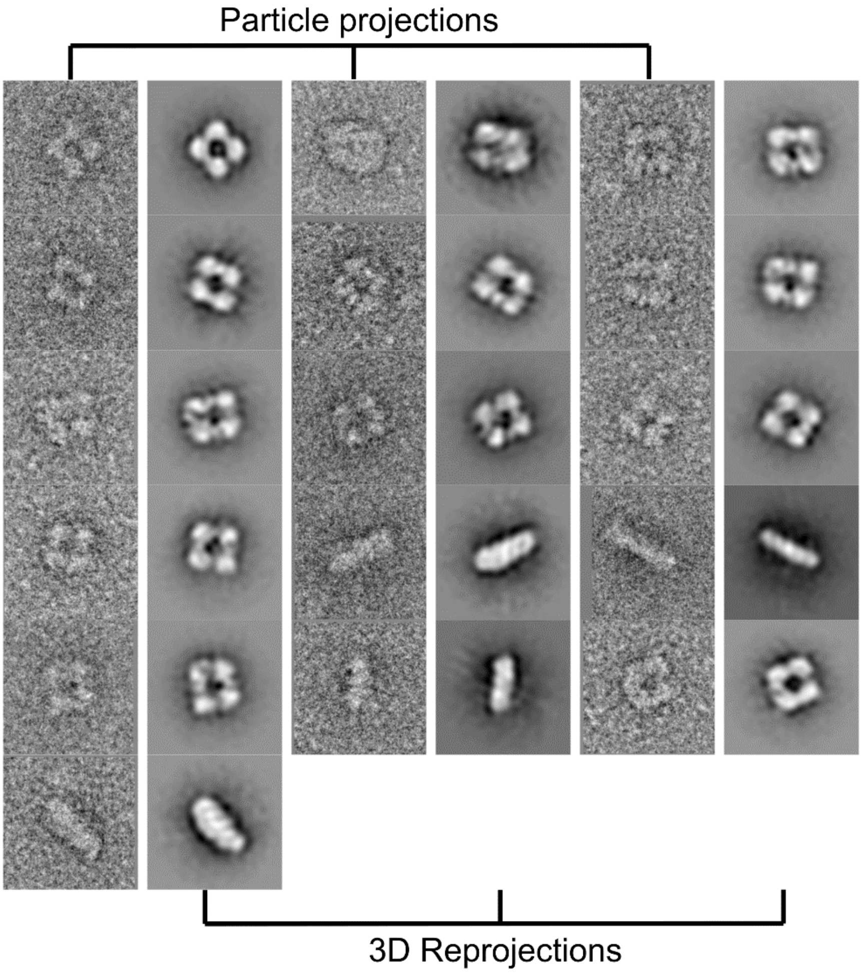


**Figure S6.** Comparison between square-like particles projections (even columns) and their reprojections from the 3D density map obtained in Relion (odd columns) to validate the 3D reconstruction.

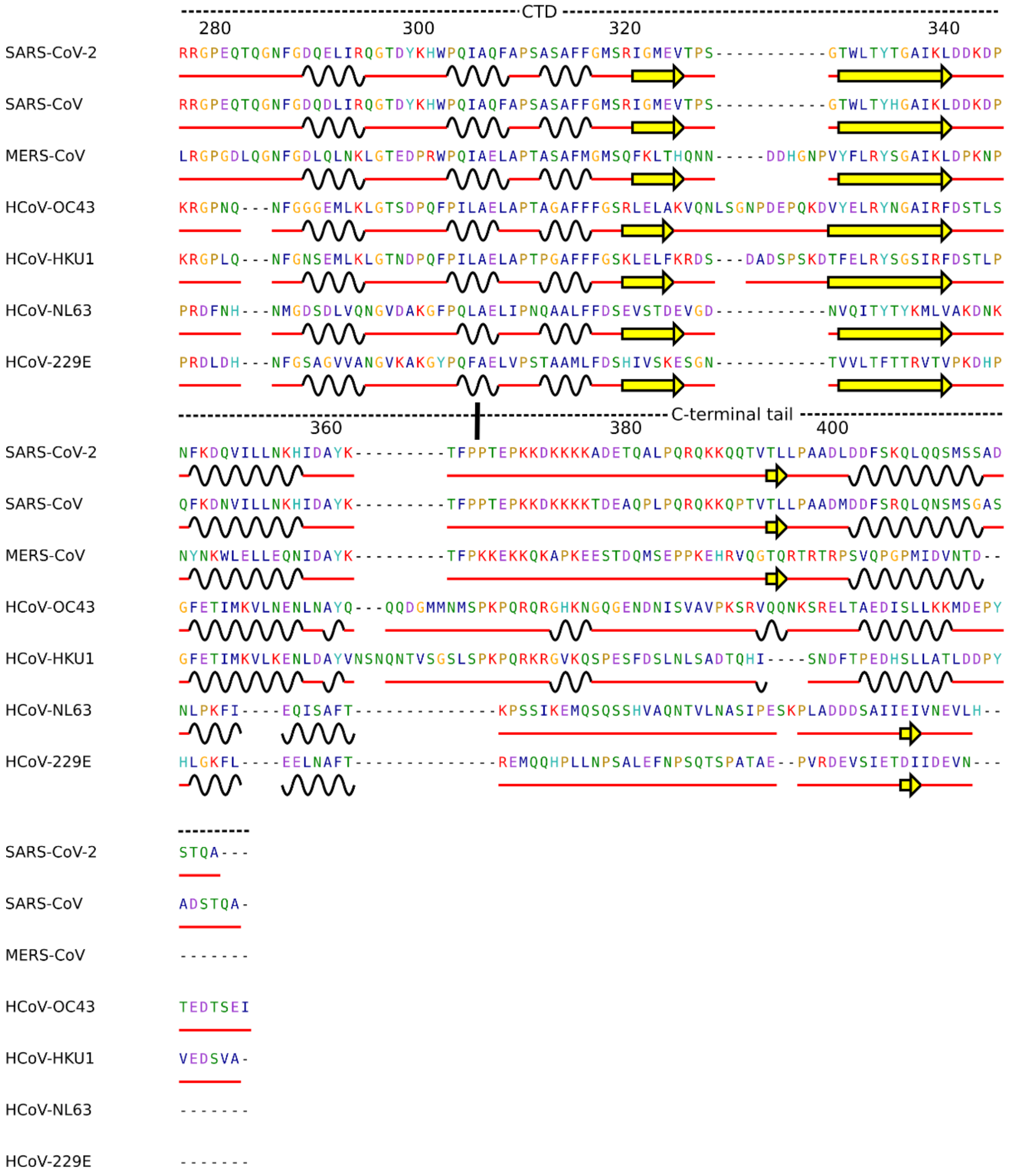


**Figure S7.** Protein sequence alignment and secondary structure prediction of the C-terminal region of coronavirus N proteins. Multiple sequence alignment was performed using Muscle algorithm (3.8) in EMBL-EBI webserver[2]. The alignment was to submitted to Ali2D webserver[3,4] and secondary structure prediction for each sequence was generated using PSIPRED algorithm. The multiple sequence alignment and secondary structure prediction was plotted using 2dSS webserver[5]. The numbers indicate SARS-CoV-2 amino acid position and does not consider the gaps. SARS-CoV-2 domains are annotated on the top of alignments.

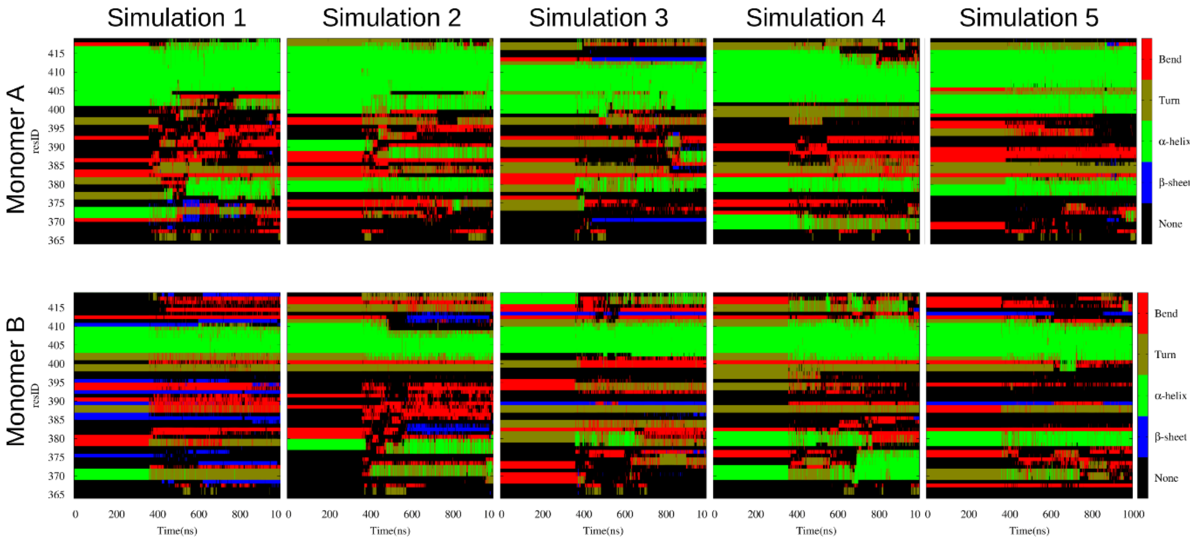


**Figure S8.** Secondary structure variation by residue for each monomer along simulation time for each of the five replicates and each monomer. The bar shows the color for each secondary structure. The figure was made using *secstruct* command from CPPTRAJ, a Ambertools20's program, and GNUPLOT.

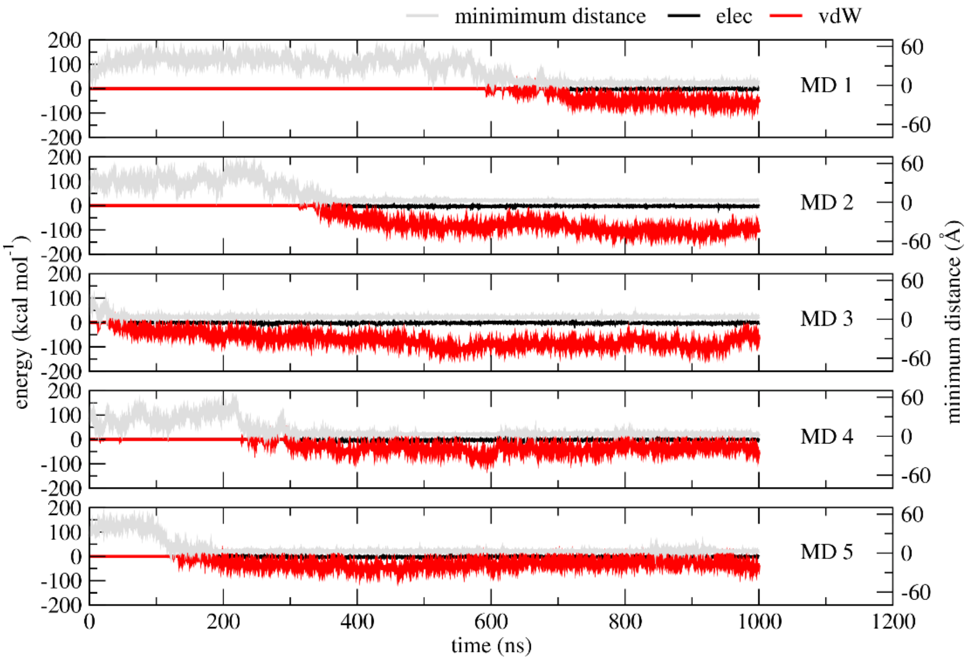


**Figure S9.** Five independent simulations of the two C-terminal tail monomers in which the dimer is spontaneously formed. The plots show the non-bonded contribution (vdw in red line and electrostatic in black line). The plots suggest the mainly non-bonded contribution is vdw.

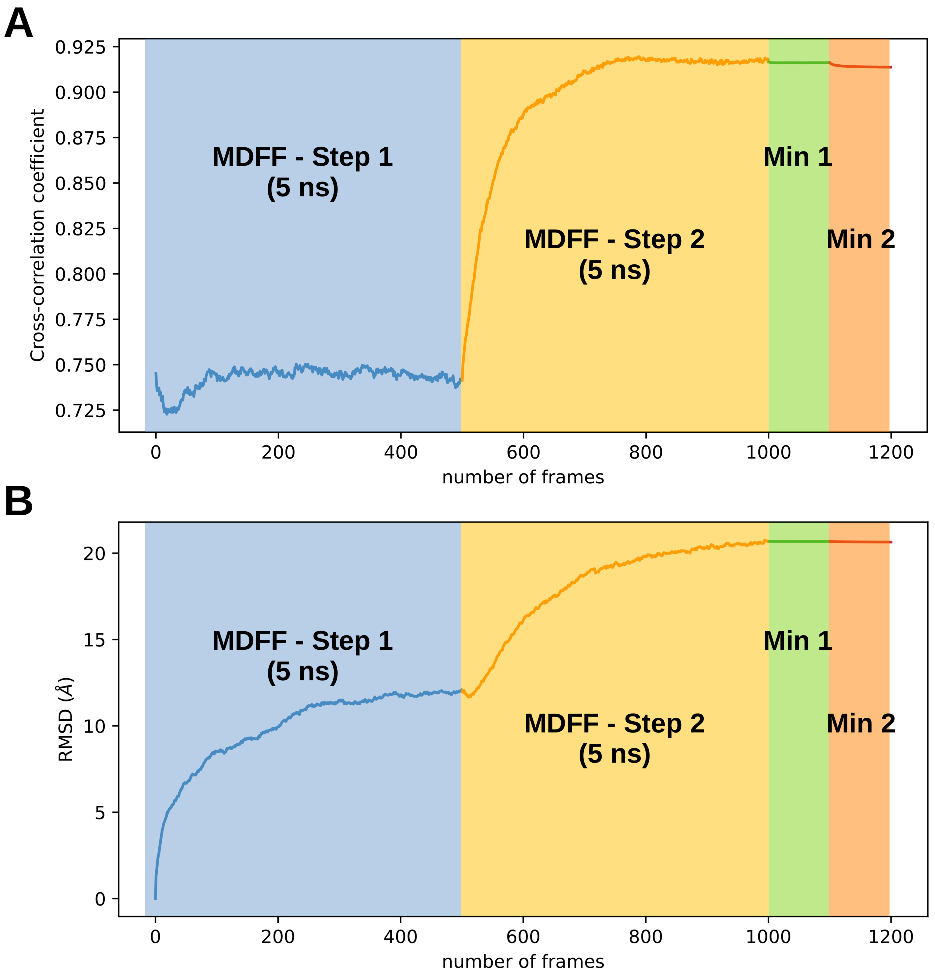


**Figure S10.** A) Cross-correlation coefficient (CCC) between model (protein and RNA) and density map along the MDFF protocol. B) RMSD of the protein and RNA backbone along the MDFF protocol. The Step 1 and Step 2 are MDFF simulations. In the Step 1, only RNA and structured domains of the N protein were included in the energy potential of the map, while in the Step 2 all protein domains and RNA were included. The Min 1 is an energy minimization step (10000 steps) with restrains on the protein backbone atom positions, while Min 2 is an energy minimization step without any restrain (10000 steps).

### References - Supplementary Material
